# Plasticity in a *Drosophila* wing circuit supports an adaptive sleep function

**DOI:** 10.1101/691451

**Authors:** K. Melnattur, B. Zhang, P.J. Shaw

## Abstract

Sleep is a near universal phenomenon whose function remains controversial. An influential theory of sleep function posits that ecological factors that place animals in harm’s way increase sleep as a state of adaptive inactivity. Here we find that manipulations that impair flight in *Drosophila* increase sleep. Further, we identify a novel neural pathway from peripheral wing sensory neurons to the central brain that mediates the change in sleep. Moreover, we show that flight impairments activate and induce structural plasticity in specific projection neurons to support increases in sleep over days. Thus, chemosensory neurons do not only signal sensory cues but also appear to provide information on wing-integrity to support behavioural adaptability. Together, these data provide mechanistic support of adaptive increases in sleep and highlight the importance of behavioural flexibility for fitness and survival.

## Main Text

Sleep has been observed in every animal that has been examined, from jellyfish to humans^1,2^. While sleep is a near universal phenomenon, animals exhibit a wide range of sleep times^3^. Interestingly, there appears to be little in the way of a discernible relationship between sleep time and cognitive capacity – sleep times can vary dramatically even among animals believed to have sophisticated cognitive abilities, chimpanzees and bonobos sleep up to 10-11 hours a day^4^ and elephants as little as 2-3 hours^5^. Further, individuals within a species also sleep to varying amounts, and sleep times vary within a given individual’s lifetime. A compact and unifying function for sleep has remained elusive perhaps due to the challenges of accounting for the diversity of sleep behaviour.

Indeed, it is increasingly clear that sleep itself is plastic, shaped by ecological factors and is responsive to environmental changes within an individual’s lifetime^6-9^. Animals from pectoral sandpipers^10^ and swallows^11^ to dolphins^12^ and elephants^5^ suppress sleep at specific times (and on occasion dispense with it altogether) without exhibiting a sleep rebound, or affecting cognition or fitness. Further, animals from *Drosophila* to humans, acutely suppress sleep in response to starvation, without suffering any apparent negative consequences^13-16^. These and other related observations have been advanced in support of an influential theory of sleep function which holds that sleep serves to maintain animals in a state of adaptive inactivity^17^ i.e. animals sleep to stay out of harm’s way, and not to serve a restorative function *per se*. An unstated assumption of the adaptive inactivity hypothesis is that inactivity is actively regulated and under the influence of natural selection. Surprisingly, neither the underlying circuitry nor the molecular mechanisms regulating sleep during dangerous or life-threatening conditions are known.

Here we show that manipulations that impair flight in *Drosophila* (and thus, the ability to stay out of harm’s way) increase sleep. The increase in sleep is signalled by chemosensory neurons that communicate with the central brain. Thus, while a primary focus of sensory biology is to elucidate how activation of a peripheral sensory sheet is transformed by the central brain to extract representations of features (e.g. edges / objects), our data indicate that this sensory information also reaches brain areas controlling motivated behaviour, notably sleep. Taken together, our data provide mechanistic evidence supporting the adaptive inactivity hypothesis and new insight into how sensory processing controls sleep need.

### Blocking wing expansion increases sleep

Newly eclosed flies emerge with unexpanded wings and in the first 20-30min post eclosion, undertake a stereotyped series of behaviours to expand their wings^18,19^. Confining flies with unexpanded wings to a restricted space overnight greatly delays wing expansion, resulting in flies that cannot fly^20^, (Fig. 1a). As seen in Fig. 1b, flies placed into confinement pre-expansion, sleep more compared to age matched siblings that were confined for the same amount of time but immediately following wing-expansion or unconfined siblings when released into recovery the following day. The increase in sleep was particularly marked during the day, and associated with increased daytime sleep consolidation (Fig. 1, b-d). Importantly, confinement did not reduce waking activity indicating locomotion was not impaired (Extended Data Fig. 1a). Further, sleep of flies that had been confined was rapidly reversible by a mechanical stimulus (Fig. 1e) and was associated with increased arousal thresholds (Fig. 1f) indicating confinement did not induce a behavioural malaise. Thus, confinement induces a state that meets the established criteria for sleep^21,22^.

**Fig. 1:**
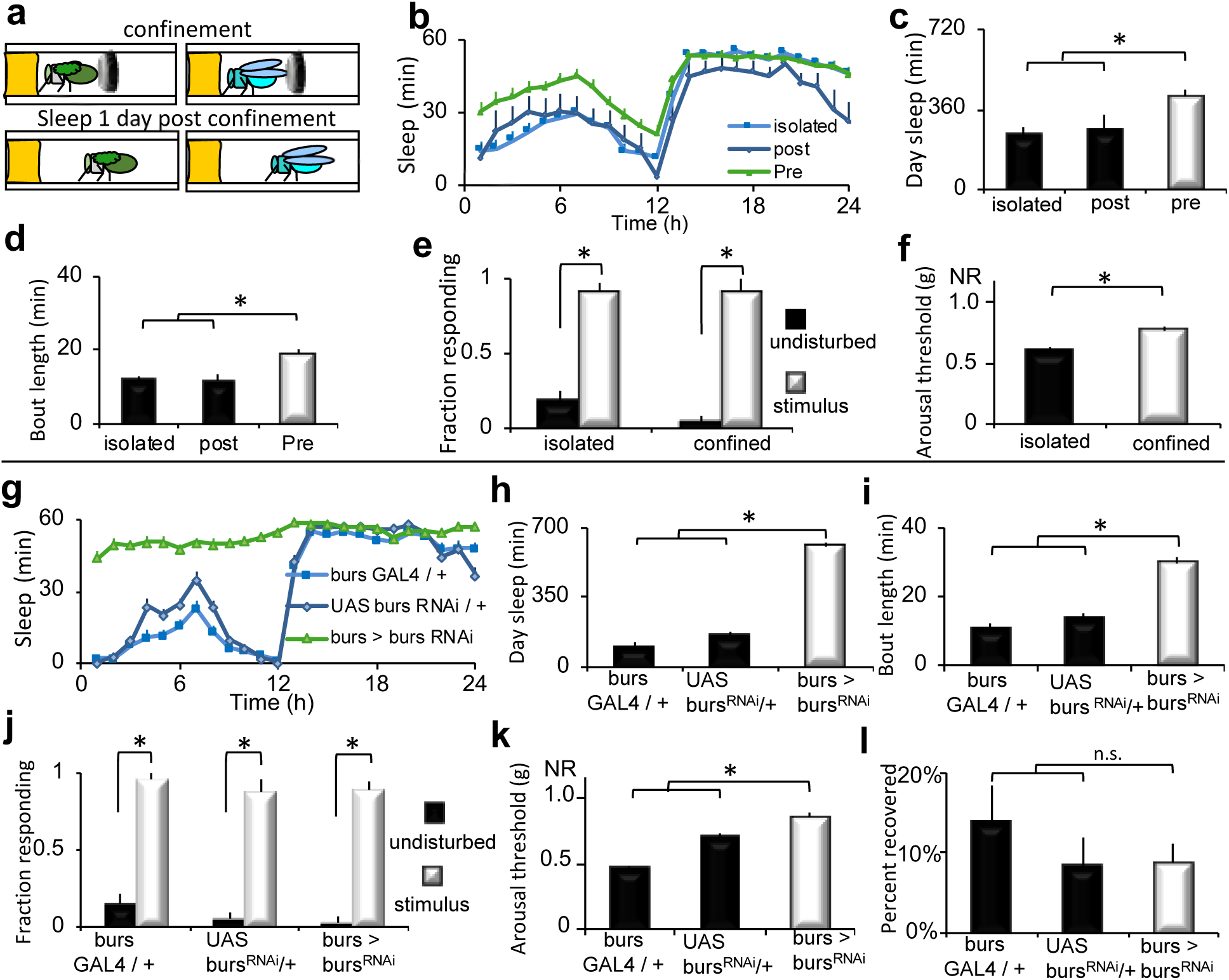
Disrupting wing expansion increases sleep. **a**, Wild-type flies were confined in a restricted space pre or post wing-expansion following eclosion (day-0) and evaluated for sleep beginning on day-1. **b**, Flies that were confined pre-expansion (pre, n=37), slept more than age-matched flies confined post-expansion (post, n=9) and unconfined, siblings (isolated, n=36); Repeated measures ANOVA Time X Condition, p< 0.05. **c**,**d**, Flies with unexpanded wings displayed increased daytime sleep and sleep bout duration compared to controls (ttest; * p=0.001). **e**, Sleep in both groups was rapidly reversible in response to a mechanical stimulus at ZT15 (n=20-32 flies per condition (*p< .01 Tukey correction). **f**, Arousal thresholds were higher in flies with unexpanded wings than isolated controls (n=14 flies per condition, * p<0.01, ttest). **g**, *bursGAL4/+>UAS-burs*^*RNAi*^*/+* flies slept more than parental controls (n=32 flies / genotype, Repeated measures ANOVA for Time X Genotype, p< 0.001). **h, I**, *bursGAL4/+>UAS-burs*^*RNAi*^*/+* displayed increased daytime sleep and sleep bout duration compared to controls (*p< .01 Tukey correction). **j**, Sleep was rapidly reversible in response to a mechanical stimulus for all genotypes (n=25-30 flies / condition, *p< .01 Tukey correction). **k**, Sleep in *bursGAL4/+>UAS-burs*^*RNAi*^*/+* flies was associated with increased arousal thresholds (n=14 flies per condition; *p< .01 Tukey correction). **l**, All genotypes displayed similar sleep rebound following 12 h of sleep deprivation (n=30-31 flies / condition).

Confinement results in unexpanded wings due to alterations in the release of the neurohormone bursicon^20^ a cystine-knot heterodimer composed of two subunits – *bursicon* (*burs*), and *partner of bursicon* (*pburs*). Loss of function of *burs* or *pburs* completely blocks wing expansion^18^. Thus, we asked whether knocking down *burs* or *pburs* would increase sleep in the absence of confinement. As seen in Fig. 1g-i, *burs*GAL4>*burs*^RNAi^ flies slept more than their parental controls, exhibiting increased daytime sleep, and sleep consolidation, without impairing locomotion (Extended Data Fig. 1b). Importantly, *bursGAL4*>*burs*^RNAi^ flies were rapidly awakened by a mechanical stimulus (Fig. 1j), exhibited elevated arousal thresholds (Fig. 1k), and a normal homeostatic response to overnight sleep deprivation (Fig. 1l). Similarly sleep amount and consolidation were also elevated in *bursGAL4*>*pburs*^RNAi^ flies without impairing locomotion (Extended Data Fig. 1, c-f). *bursGAL4>pburs*^RNAi^ flies also met the established criteria for sleep (Extended Data Fig. 1, g-i). The *burs*GAL4 and *burs*^RNAi^ flies were outcrossed to a reference wild-type strain. Thus, RNAi mediated *burs*-knockdown phenocopies the results using confinement. Flies don’t sleep well when they are confined to a small space, however *bursGAL4*>*burs*^RNAi^ flies did not lose any sleep relative to controls on the first day of adult life (Extended Data Fig. 1, j-l). Loss of function *burs* point mutations also increased sleep (Extended Data Fig. 1, m-o). Importantly, RNAi knockdown of genes that co-express with *bursicon* such as *crustacean cardioactive peptide* (CCAP), *myoinhibiting peptide precursor* (*mip*), the RNA binding protein *Lark*, or the histone transferase *absent, small, or homeotic discs 1* (*ash*) did not change sleep or affect wing expansion^23,24^ (Extended Data Fig. 1, p, q), supporting the specificity of the *burs* knockdown experiments above. Confinement and loss of *burs* function both perturb wing expansion and increase sleep.

### Two *burs*+ neurons modulate sleep

*burs* is transiently expressed in a small subset of neurons in the fly CNS - two in the subesophageal ganglion (‘Bseg’) and 12-14 neurons in the abdominal ganglion (‘Bag’) (bursGAL4>GFP, Fig. 2a). Thus, we conducted a series of experiments to determine if wing expansion and sleep could be dissociated functionally, temporally or spatially. First, *burs*+ neurons were chronically inhibited by expressing the inward rectifying potassium channel *UAS-Kir2.1*^*25*^. *bursGAL4*>*UAS-Kir2.1* disrupted wing expansion, and increased both sleep and sleep consolidation during the day (Fig. 2, b-d). The sleep episodes displayed the defining behavioural hallmarks of sleep without inhibiting locomotion (Extended Data Fig. 2, a-d). Second, we constitutively activated *bur*s+ neurons by expressing the bacterial sodium channel UAS-*NaChBac*^*26*^ with *bursGAL4*. This manipulation also disrupted wing expansion and increased sleep while meeting the behavioural criteria for sleep and without inhibiting locomotion (Extended Data Fig. 2, e-k). Similar results were obtained when the larger group of CCAP+ neurons (that includes *burs*+ neurons) were activated (Extended Data Fig. 2, l-s). Activation of CCAP neurons was shown to deplete bursicon levels in central processes^26^ suggesting a possible mechanism by which activation and inhibition yield the same phenotype. Third, we used the TARGET system^27^ to determine if *burs* GAL4 activity that supports sleep and wing expansion were temporally dissociable (Extended Data Fig. 2, t,u). Consistent with *burs* expression peaking at eclosion^26^, these experiments indicated that *burs* neuron activity was required in pharate adults/early adult life for wing expansion and sleep. Finally, we used split-GAL4 lines^28^ to specifically inactivate subsets of *burs* GAL4 expressing neurons. Inactivation of the Bseg (Bseg>UAS-Kir2.1) had a partially penetrant effect on wing expansion; the flies with wing defects increased sleep (Fig. 2, e-g, Extended Data Fig. 3, a-d), whereas inactivating the Bag did not affect wing expansion and had a modest effect on sleep (Fig. 2, h-j, Extended Data Fig. 3e). The involvement of neurons in the SEG was confirmed using the Flipase-induced intersectional GAL80/GAL4 repression (FINGR) method to disrupt subsets of CCAP+ neurons (Extended Data Fig. 3, f-l)^29^. Collectively, these results suggest that modulating the activity of just two *burs*+ neurons in a restricted time window perturbs wing expansion and increases sleep.

**Fig. 2:**
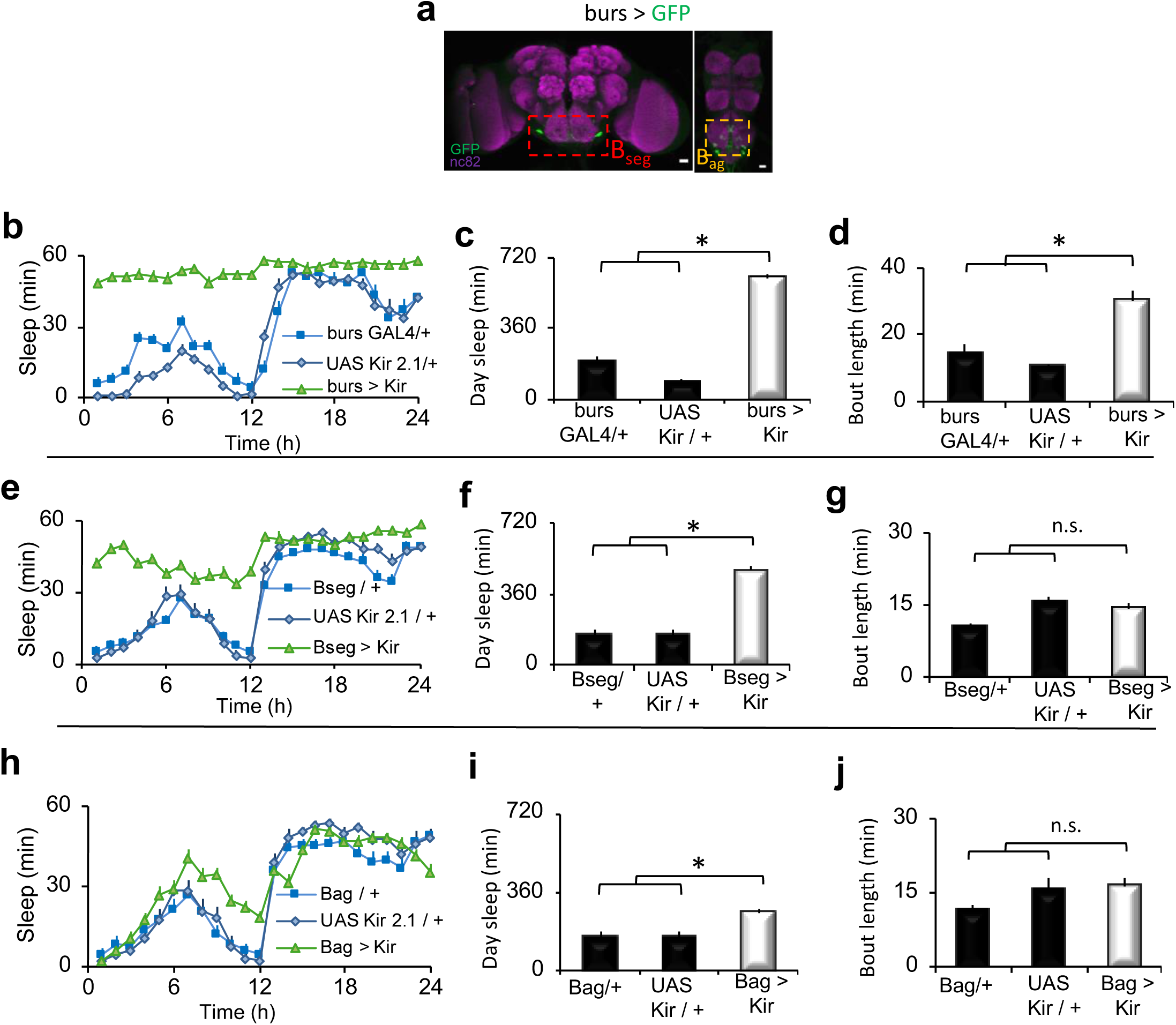
The excitability of *burs* neurons regulates wing expansion and sleep. **a**, bursGAL4/+>UAS-GFP/+ (green fluorescent protein) labels two neurons in the subesophageal ganglion (B_seg_), and 12-14 neurons in the abdominal ganglion (B_ag_). Scale bar 20μm. **b**, *bursGAL4/+>UAS-Kir2.1/+*flies slept more than parental controls (n=20-27 flies / genotype. Repeated measures ANOVA for Time X Genotype, p< 0.001) **c**,**d**, *bursGAL4/+>UAS-Kir2.1/+* flies displayed increased daytime sleep and sleep bout duration compared to controls (*p< .01 Tukey correction). **e**, *B*_*seg*_*>UAS-Kir2.1* flies with unexpanded wings slept more than parental controls (n=16-32/genotype; Repeated measures ANOVA for Time X Genotype, p< 0.001). **f, g**, *B*_*seg*_*>UAS-Kir2.1* had more daytime sleep but sleep consolidation was not altered (*p< .01 Tukey correction). **h**, *B*_*ag*_*>UAS-Kir2.1* flies had normal wings and modestly increased sleep (n=18-30 flies/ genotype Repeated measures ANOVA for Time X Genotype, p< 0.001). **i, j**, *B*_*ag*_*>UAS-Kir2.1* flies had more daytime sleep but sleep consolidation was not altered (*p< .01 Tukey correction).

### Defining the role of the bursicon receptor *rk*

The *burs* receptor is *rickets* (*rk*). rk is a leucine-rich repeat containing G-protein coupled receptor (GPCR) that signals through adenyl cyclase and protein-Kinase A (PKA)^30,31^. The strong effects we observed with manipulating *burs* function, led us to explore the effects of manipulating *rk* signalling using *rkGAL4* (Fig. 3a)^32^. We first knocked down *rk* with RNAi in *rk*+ neurons. This manipulation had a partially penetrant effect on wing expansion. *rkGAL4>rk*^RNAi^ flies with wing expansion defects exhibited increases in daytime sleep amount and consolidation without impairing locomotion (Fig. 3, b-d, Extended Data Fig. 4, a-d). *rkGAL4>rk*^RNAi^ flies with expanded wings had a small increase in daytime sleep (Extended Data Fig. 4, a-d). Point mutants in *rk* also increased sleep (Extended Data Fig. 4, e-g). We next blocked PKA signalling in *rk*+ neurons, by expressing a *dominant negative PKA* (*PKA-DN*)^33^ with *rkGAL4*. This manipulation blocked wing expansion and increased sleep amount and consolidation compared to parental controls, without impairing locomotion (Fig. 3, e-g, and Extended Data Fig. 4h). Similar results were obtained when cAMP levels were reduced in *rk*+ neurons by overexpressing the cAMP phosphodiesterase *dunce* (Extended Data Fig. 4, i-k). When is *rk* required for wing expansion and sleep? To address this question, we transiently inactivated *rk*+ neurons by expressing the temperature sensitive dynamin *shibire* (*Shi*^ts^) with *rkGAL4*. Transient inactivation for 1.5 hr post eclosion (and pre wing-expansion) was sufficient to block wing expansion and increase sleep (Extended Data Fig. 4, l-r). Collectively, these data indicate that loss of *rk* function phenocopies the effects seen with loss of *burs* function. We next examined the effects of activation of *rk*+ neurons. Chronic activation of *rk*+ neurons with *NaChBac* blocked wing expansion and increased sleep (Extended Data Fig. 5, a-d). *burs* labels 14-16 cells that die by apoptosis in the first 48hrs post eclosion^34,35^. *rkGAL4* in contrast, labels a large number of cells which persist throughout adult life. Can *rk*+ neurons regulate sleep in older flies? To address this, we activated *rk*+ neurons by expressing the heat-sensitive *transient receptor potential* 1 (*UAS-TrpA1*) channel with *rkGAL4* in 4-5 day old flies. Transient TRPA1 mediated activation of *rk*+ neurons increased sleep (Extended Data Fig. 5, e - g), indicating that *rk*+ neurons are sleep-promoting. Precisely how *rk+* neurons can increase sleep in response to such a diverse number of genetic manipulations is unclear and is the subject of ongoing investigations.

**Fig. 3:**
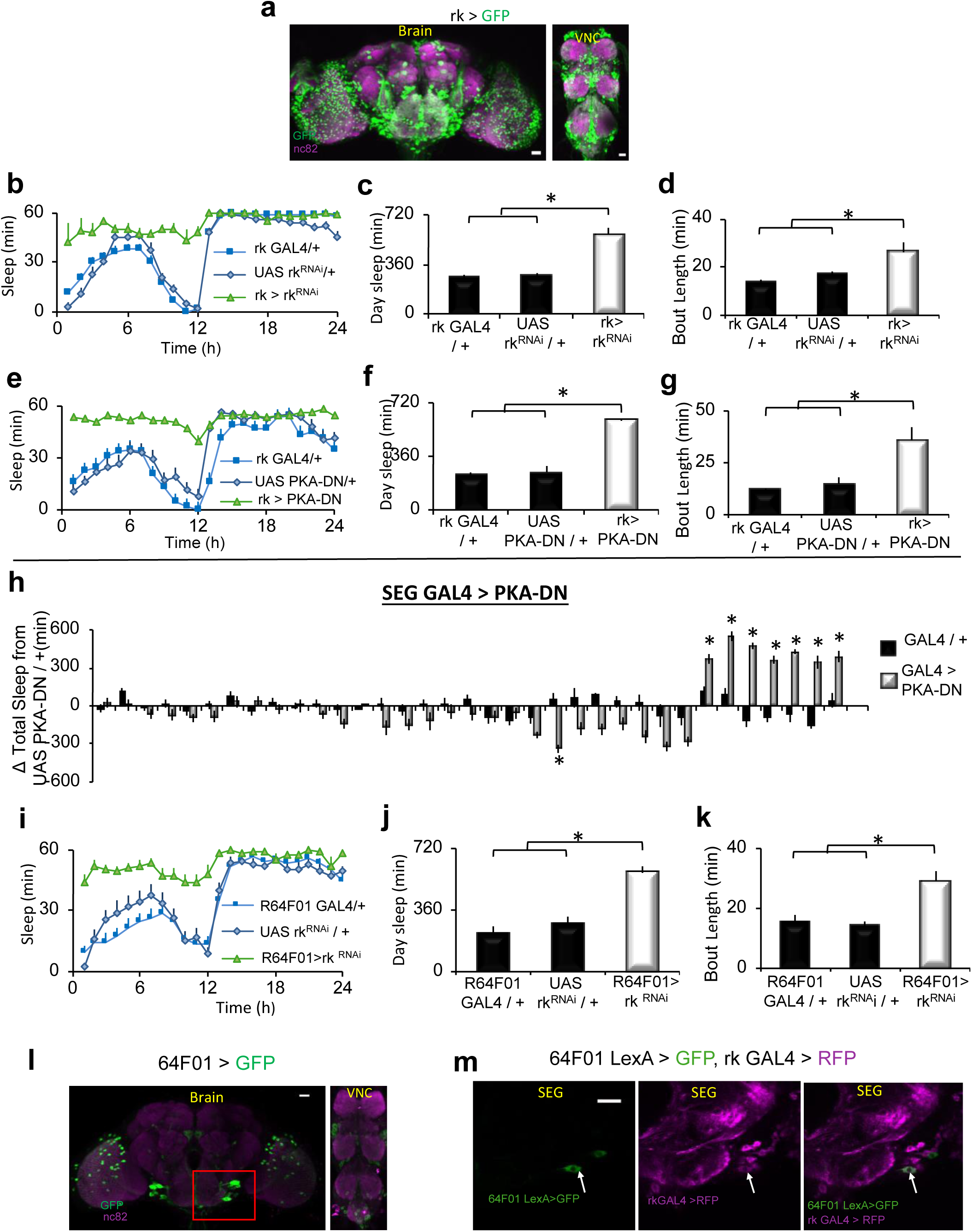
*rickets* regulates wing expansion and sleep. **a**, *rk*GAL4/+>UAS-GFP/+ labels a large number of cells in the fly CNS. **b**,**e**, Sleep was increased in both *rkGAL4/+>UAS-rk*^*RNAi*^*/+* and *rkGAL4/+>UAS-PKA^DN^/+* flies compared to parental controls (n=16-32 flies/ genotype, Repeated measures ANOVA for Time X Genotype, p< 0.001). *rkGAL4/+>UAS-rk*^*RNAi*^*/+* and *rkGAL4/+>UAS-PKA^DN^/+* displayed increased daytime sleep (**c**,**f**) and sleep bout duration (**d**,**g**) compared to controls (*p< .01 Tukey correction). **h**, Screen for SEG GAL4 drivers that increase sleep when expressing *UAS-PKA^DN^*; sleep is expressed as change in sleep in min relative to the *UAS-PKA^DN^/+* controls (*p< .01 Tukey correction). **i**, *64F01GAL4/+>UAS-rk*^*RNAi*^ flies with unexpanded wings slept more than parental controls (n=16-32 flies/genotype; Repeated measures ANOVA for Time X Genotype, p< 0.001). **j-k**, *64F01GAL4/+>UAS-rk*^*RNAi*^ flies displayed increased daytime sleep and sleep bout duration compared to controls (*p< .01 Tukey correction). **l**, *R64F01GAL4/+>UAS-GFP/+* labels a sparse population of cells in the CNS, including the SEG (red box). **m**, *R64F01LexA/+>LexAopGFP/+; rkGAL4/+>UAS-RFP/+* (red fluorescent protein) overlap in one cell in the SEG (white arrow); single confocal slices. **a, l**, Maximal intensity z-projections counterstained with nc82 (magenta). Scale bar - 20μm.

Where is *rk* required? We focused on two candidate regions – the *pars intercerebralis* (PI) a known sleep regulatory centre (where we see *rkGAL4* expression), and the SEG (as our results above implicated the Bseg). We anatomically selected a panel of GAL4 lines that label subsets of PI and SEG cells. Blocking PKA in the PI did not increase sleep (Extended Data Fig. 5h). In contrast, 8 GAL4 lines (selected from the large Rubin collection^36^) that express in the SEG increase sleep when expressing PKA-DN (Fig. 3h). To verify specificity for *rk*, we knocked down *rk* with RNAi using the primary screen hits (Extended Data Fig. 5i). Seven out of the eight lines increased sleep and blocked wing expansion in this secondary screen. We focused on one of the hits - *R64F01GAL4.* This line is derived from enhancer elements of the CCAP receptor (CCAPR) gene and since *burs* & CCAP are co-expressed, we reasoned that *R64F01GAL4* might express in a subset of rk+ neurons. We first evaluated this possibility with functional experiments. Expressing *rk*^RNAi^ (Fig. 3, i-k, Extended Data Fig. 6, a-d) or *Gβ*13F^RNAi^ (a known component of *rk* signalling) (Extended Data Fig. 6, e-g) with *R64F01GAL4* blocked wing expansion and increased sleep. In addition, expression of UAS*-NaChBac* (Extended Data Fig. 6, h-k) or *UAS-TrpA1* (Extended Data Fig. 6, l-n) with *R64F01GAL4* increased sleep, similar phenotypes to those obtained with *rk*GAL4. *R64F01GAL4* displays a restricted expression in the fly CNS, enriched in the SEG (Fig. 3l). Although *R64F01LexA* does not fully recapitulate the R64F01-GAL4 expression pattern, expressing LexAop-GFP using R64F01-LexA and UAS-RFP using *rk*GAL4 identifies at least one common neuron in the SEG (Fig. 3m).

Finally, to identify a minimal subset of R64F01 neurons that mediate the effects on wing expansion and sleep, we used an intersectional approach where we combined GAL80s with *R64F01GAL4>rk*^RNAi^. *dvGlut GAL80;R64F01GAL4>rk*^RNAi^ flies had normal (expanded) wings and unchanged sleep (Extended Data Fig. 7, a-d), suggesting that it is the glutamatergic *R64F01GAL4* neurons which are the critical subset for wing expansion and sleep. Indeed, most *R64F01GAL4* neurons appear to be glutamatergic (Extended Data Fig. 7e).

### Disrupting wings increases sleep

Wing damage occurs in adults to negatively impact flight^37^. To determine whether the adaptive inactivity role for sleep would be observed in adults, we cut wings of flies on the first day of adult life after they had expanded their wings and examined sleep two days later. Flies with wings cut increased both sleep and sleep consolidation during the day (Fig. 4, a-c) and night (Extended Data Fig. 7, f, g) compared to their siblings with intact wings; locomotion was not impaired (Extended Data Fig. 7h). Sleep of flies with cut wings was rapidly reversible (Extended Data Fig. 7i), and associated with increased daytime arousal thresholds (Extended Data Fig. 7j). To determine whether flies with cut wings were under higher sleep drive, we expressed the calcium dependent nuclear import of LexA (CaLexA) system to monitor activity of the ellipsoid body R2 neurons (a known marker of sleep drive^38^). Wing-cut increased CaLexA signal in the R2 neurons suggesting that sleep drive was increased (Fig. 4, d, e). Importantly, the wing-cut mediated sleep-increase was not a response to wing damage/injury as mutations in immune response genes did not impair the ability of wing cut to induce sleep (Extended Data Fig. 7k)^39^. Further, increases in sleep were also observed when wings were glued (Extended Data Fig. 7, l-o). Similarly,). Finally, we evaluated a number of genetic manipulations that impair flight. Expressing the cell death activator *reaper (UAS-rpr)* in the wing disc, or mutations in *wingless (wg*), protein kinase c (pkcΔ), and the commonly used *CyO* marker, all impair flight^40,41^ and increase sleep (Extended Data Fig. 8, a-e). Since many of these mutations are likely to have pleiotropic effects, we focused on wing-cut.

**Fig. 4:**
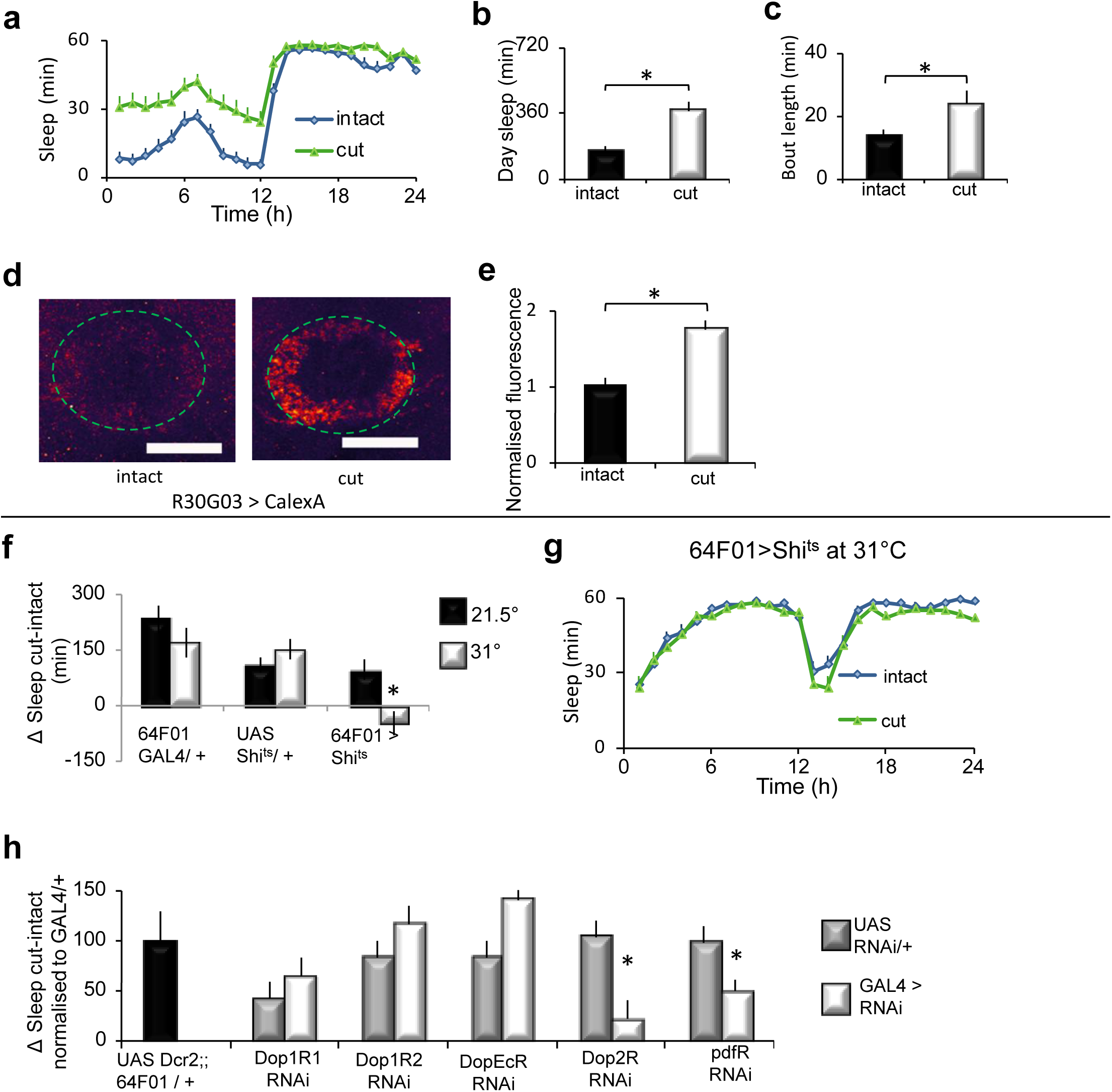
Cutting wings increases sleep. **a**, Flies slept more the second day following wing-cut than age-matched controls (n=32 flies / condition); Repeated measures ANOVA Time X Condition, p< 0.001. **b**,**c**, Flies with cut wings displayed increased daytime sleep and sleep bout duration compared to controls (ttest; * p=0.001). **d**, Representative images of R30G03/+>CalexA/+ with intact and cut wings. **e**, Quantification of CaLexA; signal was higher in brains of flies with cut wings compared controls (n=8-9 flies/condition; *p<0.001 ttest). **f**, Silencing *R64F01GAL4/+>UAS-shi*^*ts1*^*/+* neurons by raising the temperature from 21.5 °C to 31°C blocked the increase in sleep following wing cut. Sleep is expressed as a change in sleep in cut vs intact siblings. (n=16 flies per genotype / condition; *p< .01 Tukey correction). **g**, Sleep in min/hr for *R64F01GAL4/+>UAS-shi*^*ts1*^*/+* flies shown in (**f**). **h**, *R64F01GAL4/+>UAS-Dop2R*^*RNAi*^*/+* and *R64F01GAL4/+>UAS-Pdfr*^*RNAi*^*/+* flies displayed an attenuated change in sleep following wing cut; *R64F01GAL4/+>UAS-Dop1R1*^*RNAi*^*/+, R64F01GAL4/+>UAS-Dop1R2*^*RNAi*^*/+* and *R64F01GAL4/+>UAS-DopEcr*^*RNAi*^*/+* were not different from controls (n=16-32 flies / genotype; (*p< .05 Tukey correction). **d**, Maximum intensity confocal projections. Scale bar 20μm.

Disrupting wings in adult flies increases sleep. Transient activation of wing-expansion circuits in adults also increases sleep (see above). Together, these results suggest that wing-cut may recruit wing expansion circuits to increase sleep. To test this hypothesis, we expressed a thermosensitive mutant form of *dynamin, Shibire* (UAS-shi^ts1^) to block clathrin mediated endocytosis of neurotransmitter release only at non-permissive temperatures (31°C). As seen in Fig. 4, f, g at 31°C, wing-cut did not result in an increase in sleep in *64F01GAL4>UAS-shi*^*ts1*^ flies compared to either siblings maintained at 21.5°C or parental controls. Although bursicon neurons undergo apoptosis^34,35^, both dopamine and *Pigment dispersing factor* (*Pdf*) are known to regulate flight and sleep in adulthood^42-47^. Interestingly, knocking down the *D2 dopamine receptor* (*Dop2R*), the *Pdf receptor* (*Pdfr*), and their downstream signalling components in *R64F01* neurons mitigated the increased sleep following wing cut (Fig. 4h, Extended Data Fig. 8, f-h). Knocking down the other known *Drosophila* dopamine receptors did not appear to influence the extent of wing-cut induced sleep, although the precise role of the *Dop1R1* receptor remains ambiguous, and will be revisited later (Fig. 4h). Further, dopaminergic neural processes were detected in close proximity to the R64F01GAL4 neurons in the SEG (Extended Data Fig. 8i), suggesting that the SEG is the relevant site of dopaminergic modulation. Although it is possible that distinct subsets of neurons mediate wing expansion and the response to wing-cut, we feel this scenario is unlikely because the wing-cut response was normal in *dvGlutGAL80; 64F01GAL4 >Dop2R*^*RNAi*^ flies (Extended Data Fig. 8j). Impairing flight by cutting wings increases sleep, requires R64F01 neurons, and is not immune-mediated.

### A neural pathway for wing-cut induced sleep

We hypothesised that a neural pathway from the wing conveys information about wing integrity to the brain to modulate sleep. To test this hypothesis, we inactivated subsets of wing chemosensory and mechanosensory neurons using UAS-Kir2.1. Expressing UAS-Kir2.1 with either *Ir52aGAL4* or *Ir76bGAL4* attenuated but did not block the increase in sleep following wing-cut (Fig. 5a). These data suggest that the combined input from *Ir52aGAL4* and *Ir76bGAL4* is additive such that the smaller increase in sleep may result from losing input from the non-silenced set of neurons during wing cut. Indeed, *Ir52aGAL4>dTrpA*1 flies sleep more than parental controls confirming that these neurons can modulate sleep (data not shown). *Ir52a GAL4* strongly expresses in putative chemosensory neurons along the wing margin that project into the wing neuromere of the ventral nerve cord (VNC), is weakly expressed in leg sensory neurons that also project axons into the VNC but not detected in the brain (Fig. 5b)^48^. *Ir76b* GAL4 expression is similar to *Ir52a* GAL4, and in addition, expresses in other classes of sensory neurons (Extended Data Fig. 9a). We focused on *Ir52a* GAL4, as its expression pattern was more restricted.

**Fig. 5:**
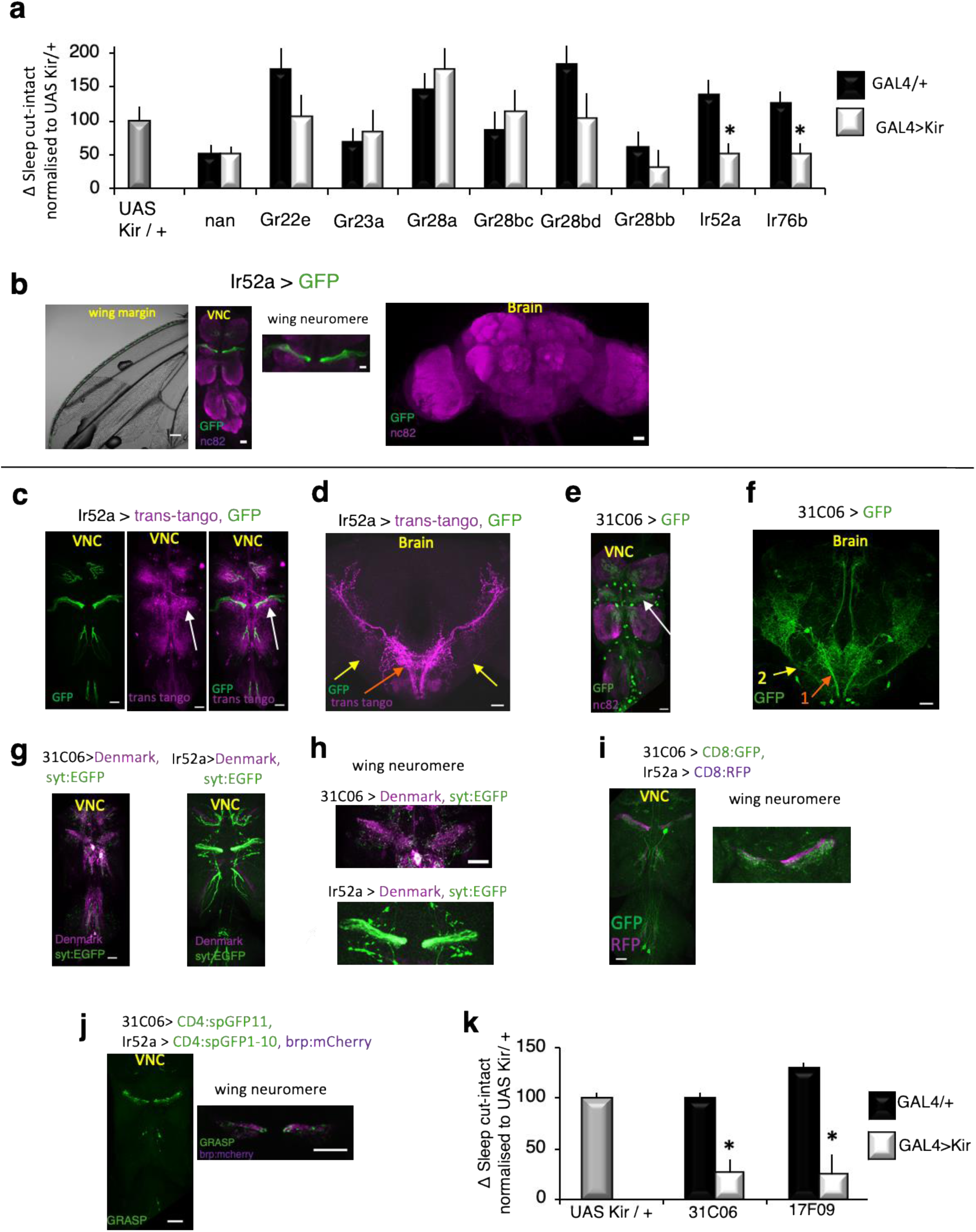
Circuit for wing-cut induced sleep. **a**, Change in sleep (cut-intact) in flies expressing *UAS-Kir2.1* in GAL4 lines associated with wing chemo and mechano-sensation normalised to the UAS-Kir2.1 parental control. *Ir52aGAL4/+>UAS-Kir2.1/+* and *Ir76bGAL4/+>UAS-Kir2.1/+* flies did not increase sleep in response to wing cut (n=20-45 flies /condition; * p <0.05, Tukey correction). **b**, Ir52aGAL4/+>UAS-GFP/+ labels subsets of wing neurons (left) that project into the wing neuromere of the VNC (middle). Weak expression was also detected in nerves from leg neurons that project into the VNC (middle). No expression was detected in the brain (right) **c**, *Ir52aGAL4/+>Trans-tango/+* (magenta) detects neurites in close proximity to the projections of *Ir52aGAL4* axons (green) in the VNC, with prominent labeling in the wing neuromere (white arrow), and two projection neuron axon tracts that exit the VNC and project to the lateral protocerebrum. **d**, *Ir52aGAL4/+>Trans-tango/+* labels VNC neurons that project axons out of the VNC into the brain in two tracts with arborisations in the SEG and the VLP (orange and yellow arrows) (**e**) In the VNC, *31C06GAL4/+>UAS-GFP* labels neurites that resemble the Ir52a>trans-tango pattern in (**c**) with strong labeling in the wing neuromere (white arrow). **f**, In the brain, *31C06GAL4/+>UAS-GFP/+* labels neurons that project in patterns similar to the Ir52a>trans-tango labeled axons (orange and yellow arrows, “1” and “2”). **g**,**h**, *31C06GAL4/+>UAS-Denmark,UAS syt_EGFP/+* and *Ir52aGAL4/+>UAS-Denmark*,,*UAS syt_EGFP/+* staining patterns. *UAS-Denmark* (magenta) labels dendrites, syt:EGFP (green) labels presynaptic sites. **i**, 31C06LexA/+>LexAop CD8:GFP/+, and *Ir52aGAL4/+>UAS CD8:RFP/+* expression patterns reveal that 31C06 dendrites (GFP, green) are in close proximity to Ir52aGAL4 axons (RFP, red), particularly in the wing neuromere (right). **j**, Strong GRASP signal was detected between *31C06LexA* dendrites and *Ir52aGAL4* axons in the VNC (left). GRASP signal was in close proximity to *Ir52aGAL4* pre-synaptic sites (right, brp:mcherry in magenta). **k**, *31C06GAL4/+>UAS-Kir2.1* and 17F09GAL4/+>UAS-Kir2.1 blocked the increase in sleep following wing cut compared to parental controls (n=27-46 flies /condition; *p< .01 Tukey correction). **b**-**j**, Maximum intensity confocal projections. Scale bar 20μm. GRASP – GFP Reconstitution Across Synaptic Partners. VNC– Ventral Nerve Cord. VLP - Ventro Lateral Protocerebrum.

What circuits are downstream of *Ir52a*+ neurons? Second order neurons to wing sensory neurons are hitherto unknown. To identify post-synaptic partners of *Ir52aGAL4* neurons, we used the *trans*-*tango* system which labels neurons one synapse from a given pre-synaptic neuron^49^. *Ir52aGAL4*>*trans-tango* labelled two classes of projection neurons with neurites in close proximity to *Ir52*a sensory axons in the VNC, particularly in the wing neuromere (Fig. 5c). These axon tracts exit the VNC, arborise in the SEG and terminate in the ventro-lateral protocerebrum (VLP) in the brain (Fig. 5d). In analogy to olfactory projection neurons, we call these tracts the medial and lateral VNC-VLP tract. *Ir76bGAL4*>*trans-tango* labelled a broader neural population including a tract that resembled the medial VNC-VLP tract above (Extended Data Fig. 9, b-d). From a visual screen of images of GAL4 lines^36^, we identified one line -*31C06 GAL4* whose expression pattern resembles the *Ir52a*>*trans-tango* pattern (Fig. 5, e, f, Extended Data Fig. 9e). *31C06GAL4* projection neurons are likely post-synaptic to *Ir52aGAL4* sensory neurons. *31C06GAL4* neurites are largely dendritic, and *Ir52aGAL4* neurons largely axonal in the VNC (Fig. 5, g, h). The processes of *31C06LexA* projection neurons and *Ir52aGAL4* sensory neurons are in close proximity (Fig. 5i), and make physical contacts that appear to be synaptic (as evidenced by GFP Reconstitution Across Synaptic Partners, GRASP, signal) (Fig. 5j). Finally, a recently developed tool to identify enhancers that overlap with a given line^50^, identified a second GAL4 line - *17F09 GAL4* that overlaps with *31C06 GAL4* in projection neurons of the VNC-VLP tract. Inactivation of VNC-VLP projection neurons with *31C06 GAL4* or *17F09 GAL4* abrogated the wing-cut response (Fig. 5k).

### Flight impairments activate and induce plasticity in projection neurons

These results describe a neural pathway that connects the wings to higher brain centres and is required for wing-cut induced sleep. Further, they suggest the possibility that wing-cut or flight-impairment more generally, directly modulates these projection neurons to increase sleep. Indeed, we found that activation of *31C06GAL4* neurons with *dTRPA1* increased sleep (Extended Data Fig. 10, a-c) and that wing-cut elevated CaLexA signal in *31C06GAL4* VNC projection neurons indicating that wing-cut activates this pathway (Fig. 6, a-c). What are the mechanisms that support the changes in activity? We hypothesised that flight-impairments would induce plastic changes in the number of synapses in *31C06GAL4* projection neurons to stably modulate sleep. To test this possibility, we used the Synaptic Tagging with Recombination (STaR) system^51^ which labels active zones in neurons of interest via recombinase based tagging of the active zone protein Bruchpilot (BRP)(Fig. 6, d-g). We observed more BRP puncta and increased BRP intensity per punctum, along the medial and lateral VNC-VLP tracts referenced above, of *31C06GAL4>STaR* flies with curly or cut wings relative to controls with normal wings (Fig. 6, h-k). In addition, we also observed increased BRP intensity per punctum in the terminal arborisations of the *31C06GAL4* projection neurons (Extended Data Fig. 10, d, e). Finally, *31C06LexA* projection neurons are pre-synaptic to the *64F01GAL4* wing-expansion/wing-cut neurons–their processes are in close proximity (Fig. 6l), exhibit complimentary axonal and dendritic profiles (Extended Data Fig. 10, f, g), and they make physical contacts that appear to be synaptic (Fig. 6, m, n). Thus, while 31C06 neurons project to several brain areas that may influence sleep, *64F01GAL4* neurons likely represent one sleep promoting output of *31C06* neurons.

**Fig. 6:**
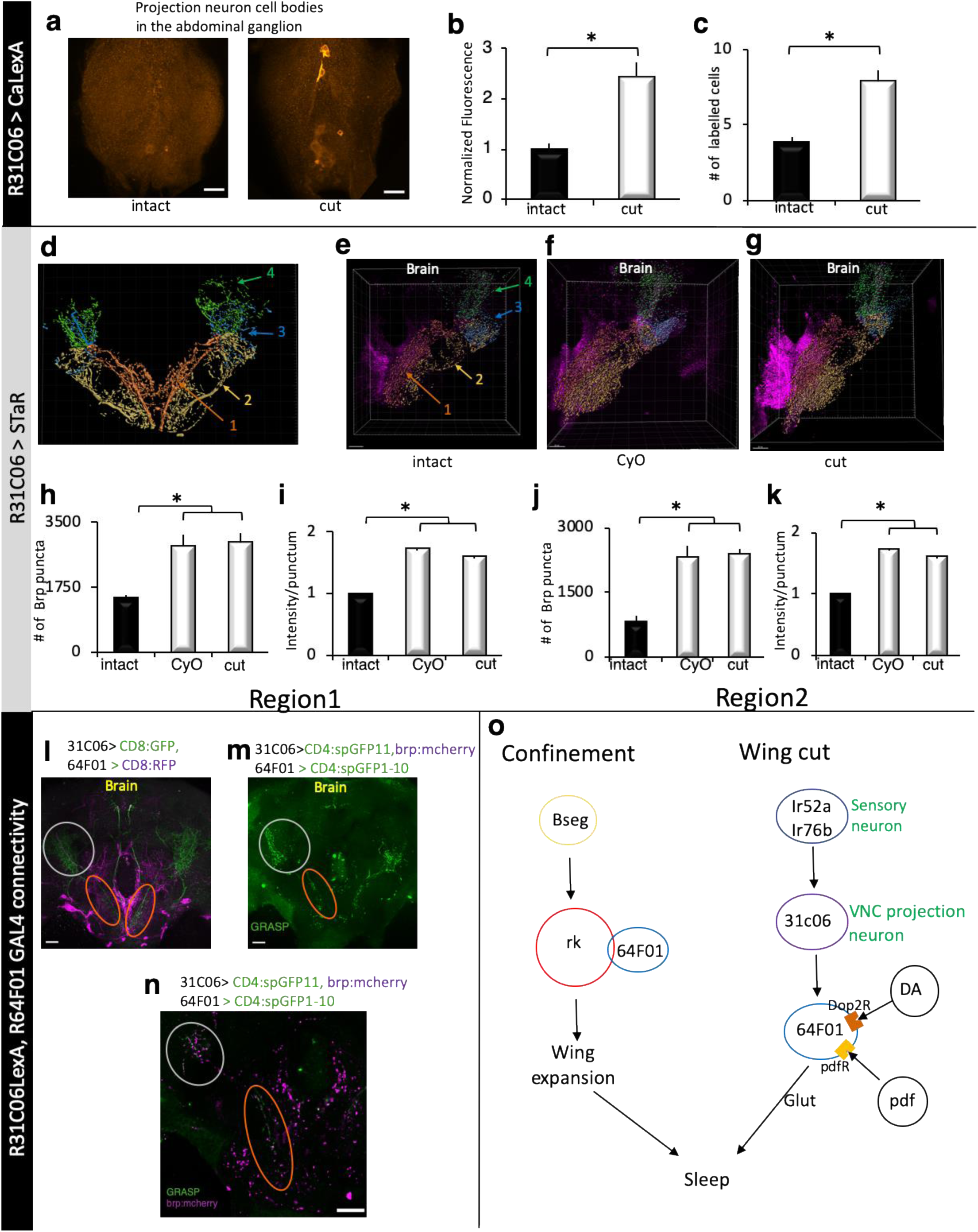
Wing cut induces structural plasticity in *31C06GAL4* projection neurons. **a**, Pseudo-coloured representative images of *31C06GAL4/+>CalexA* expression in the abdominal ganglion (AG). Wing cut increases the number (**b**) and intensity (**c**) of labelled cells compared to intact siblings (n=8-10 flies/condition; *p<0.01, and *p<0.01 ttest, respectively). **d**, Schematic of projections from R31C06GAL4; region 1 (orange-red) captures the medial projection, region 2 (gold) reflects the lateral projection, regions 3 and 4 reflect the lateral and dorsal VLP arborisations of the two projections (see methods for details). **e**-**g**, Representative images of BRP puncta in intact (**e**) and cut (**f**) *R31C06GAL4/+>STaR* flies as well as in *R31C06GAL4/+>CyO/+;STaR* flies (**g**). **h, I**, In region 1, the number and intensity of BRP puncta was increased in cut *R31C06GAL4/+>STaR* flies and *R31C06GAL4/+>CyO/+;STaR* flies compared to intact controls. **j, k**, The number and intensity of BRP puncta were also increased in region 2 in cut *R31C06GAL4/+>STaR* flies and *R31C06GAL4/+>CyO/+;STaR* flies compared to intact controls (n=7/group, *p <0.002, Tukey correction). **l**, *R64F01GAL4/+>UAS-RFP/+* (magenta), and *31C06LexA+>LexAop-GFP/+* (green) neurites are in close proximity in the SEG (orange ellipse) and the VLP (grey circle). **m**, GRASP signal (green) was detected in these regions (orange and grey circles). **n**, GRASP signal was adjacent to 31C06LexA presynaptic sites (brp:mcherry, magenta) in SEG and the VLP. **o**, Schematic of identified flight and sleep regulating circuitry DA – dopamine, pdf – pigment dispersing factor, Glut - Glutamate. (**a**,**l** &**m**) Maximal intensity confocal projections,(**n**) Single confocal slice of brain in (**m**) (**e-g**) Snapshots from IMARIS software. Scale bar 20μm.

## Discussion

Two independent methods of disrupting wings – confinement and wing-cut, increase sleep. Further, our data describe a novel neural pathway from the wings to the brain that underlies the effects of wing cut (Fig. 6o). Flight impairments induce structural plasticity (as measured by BRP puncta) in this circuit and are associated with increased sleep. BRP levels at individual active zones correlates with activity^52,53^. Further, changes in BRP levels and number and distribution of BRP puncta track activity-dependent plastic changes^54,55^. Structural plastic changes could thus be an elegant method to ensure persistence of activity and behavioural outcome (in this case, sleep) days after the insult (wing-cut).

Collectively, these data describe a surprising relationship between flight and sleep – impairing flight increases sleep. While the existence of such a relationship might at first seem esoteric, wing damage and flight impairments might be fairly common. Supporting this notion, a recent study found that male flies frequently inflicted wing damage in aggressive bouts^37^. The increases in sleep we observe when flight is impaired, could thus be viewed as an adaptive response, enabling flies to modify their behavioural repertoire to meet new challenges^40^. Indeed, sleep has been proposed to serve as a state of ‘adaptive inactivity’. In providing evidence of adaptive *increases* in sleep in an invertebrate our data support a key prediction of this idea and greatly extend its applicability.

## Methods

### Flies

Flies were cultured at 25°C with ∼50% relative humidity, and kept on a standard yeast, corn syrup, and agar diet, while being maintained on a 12hr light : 12hr dark cycle. Crosses with *UAS rk RNAi* were set up at 29°C as per established protocols^56^.

#### Fly Strains

*burs GAL4, rk GAL4* (rk-pan GAL4), *B*_*seg*_ *GAL4* (ET^VP16AD^-99 ⋂ burs Gal4DBD ^U6A1^), *B*_*ag*_ GAL4 (ET^VP16AD^-N9A88A ⋂ burs Gal4DBD^U6A1^), *UAS dnc* were gifts of Ben White (NIMH). *burs*^Z1091^, *burs*^Z5569^, and *bw; st* flies were gifts of C. Zuker (Columbia). *UAS PKA-DN*, a constitutively active regulatory PKA-subunit (UAS R*), was a gift of D. Kalderon (Columbia). *Sifa GAL4, kurs*^58^ *GAL4*, and *DH44*^VT^ *GAL4*^*57*^ were gifts of A. Sehgal (University of Pennsylvania). *9-30 GAL4* and *12-230 GAL4* ^58^ were gifts of F. Wolf (UC Merced). *UAS dTRPA1* (homozygous viable 2^nd^ chromosome insert) was a gift of P. Garrity (Brandeis). *UAS Kir2.1EGFP* (homozygous viable 3^rd^ chromosome insert) was a gift of R. Baines (Manchester). *UAS rk RNAi* (8930-R1) was obtained from NIG-FLY (National Institute of Genetics, Mishima, Japan). W4 (CCAP-GAL4, UAS-GluedDN, tubP>stop>GAL80, UAS mCD8GFP) flies and the enhancer-trap flipase (Et-flp) lines used in Extended Data Fig. 3 were gifts of B. Zhang (University of Missouri). *UAS Shi*^ts^ (pJFRC100-20XUAS-TTS-Shibire-ts1-p10 in attp2) was a gift of G. Rubin (Janelia Farms Research Campus). GRASP reagents^59,60^ – *UAS CD4:SpGFP*_*1-10*_, LexAop *CD4:SpGFP*_*11*_, *UAS brp:mcherry* (3^rd^ chromosome insert), *lexaop brp:mcherry*^*61*^ (3^rd^ chromosome insert) were gifts of C-H. Lee (Academia Sinica, Taiwan)

All other GAL4 and LexA lines were obtained from the Bloomington Drosophila Stock Center. The following lines were also obtained from the Bloomington Drosophila Stock Center: *rk*^1^, *rk*^4^, *wg*^1^, *pkcΔ*^e04408^, *rel*^E20^, *imd*^1^, *UAS NaChBac* (*UAS NaChBacEGFP4*), *20XUAS-IVS-mCD8GFP* (in attP2), *10XUAS-IVS-mCD8::RFP* (attP18) *13XLexAop2-mCD8::GFP* (attP8), *UAS rpr* (UAS-rpr.C^14^ on 2^nd^ chromosome), *dvGlut GAL80* (VGlut^MI04979-T3XG80.2^), *trans-tango* (UAS myrGFP.QUAS mtdTomatoHA; trans tango), *UAS Denmark, UAS syt.EGFP* (on 2^nd^ chromosome), *CaLexA* (LexAop-CD8::GFP-2A-CD8::GFP; LexAop-CD2::GFP; UASmLexA-VP16-NFAT / TM6B), *STaR* (UAS FLP, brp(FRT.stop)V5-2A-LexA-VP16 in VK00033), *UAS burs RNAi*^J02260^, *UAS pburs RNAi*^HMC04211^, *UAS CCAP RNAi*^HMJ23953^, *UAS CCAPR RNAi*^JF01338^, *UAS ash1 RNAi*^HMS00582^, *UAS lark RNAi*^JF02783^, *UAS Mip RNAi*^HMS02244^, *UAS Gβ13F RNAi*^HMS01455^, *UAS Gβ76c RNAi*^JF03127^, *UAS Gγ30a RNAi*^HMS01455^, *UAS Gα*_i_ *RNAi*^HMS01273^, *UAS Gα*_*o*_ *RNAi*^HMS01129^, *UAS Gα*_*q*_ *RNAi*^HMJ30300^, *UAS plc21c RNAi*^HMS00600^, *UAS Itpr RNAi*^HMC03351^, *UAS Stim RNAi* ^HMC03651^, *UAS Irk1 RNAi* ^HMS02480^, *UAS Irk2 RNAi*^HMS02379^, *UAS Irk3 RNAi*^JF02262^, *UAS Dop1R1 RNAi* ^HM04077^, *UAS Dop1R2 RNAi* ^HMC06293^, *UAS Dop2R RNAi*^HMC02988^, *UAS DopEcR RNAi*^JF03415^, *UAS pdfR RNAi*HMS01815

#### Genetics

*burs GAL4, rk* GAL4, *rk*^1^, *rk*^4^, *UAS burs RNAi*^J02260^, *UAS pburs RNAi*^HMC04211^, *UAS NaChBac, UAS Kir2.1, ET*^VP16AD^*- 99, burs Gal4DBD ^U6A1^*, ET^VP16AD^-N9A88A, were all outcrossed to a reference *yw* line for 5 generations. Using balancer chromosomes, a *yw*; *CyO* / *Sco* line was generated where the 1^st^, 3^rd^ and Y chromosomes were identical to a reference *yw* strain.

### Behavioural Analysis

#### Sleep

Sleep was assessed as previously described^62^. Briefly, individual virgin female flies were placed into 65mm tubes, and their locomotor activity continuously measured using the Drosophila Activity Monitoring (DAM) system (Trikinetics, Waltham, MA). Locomotor activity was binned in 1 min intervals; sleep defined as periods of inactivity of 5 min or more was computed using custom Excel scripts. In sleep plots, sleep in min/hr is displayed as a function of zeitgeber time (ZT). ZT0 represents the beginning of the fly’s subjective day (lights on), and ZT12 represents the transition from lights on to lights off.

#### Sleep Homeostasis

4-7 day old female flies were placed into 65mm tubes in DAM monitors and sleep recorded for 2 days to establish a baseline. Flies were then sleep deprived for 12 hrs during the dark phase (ZT12-ZT0) using the Sleep nullifying apparatus (SNAP) with procedures previously described^63^. For each individual fly, the difference between the sleep between the sleep time on the recovery day and baseline was calculated as the sleep gained / lost. Sleep rebound was calculated as the ratio of this sleep gained / lost to the sleep lost during baseline, expressed as a percentage.

#### Reversibility

Female flies were placed into 65mm tubes in DAM monitors. A mechanical stimulus was delivered for 10min at ZT15. Only flies that had been inactive for at least 5 min preceding the stimulus were considered for analysis. The fraction of flies aroused by this stimulus was computed for flies subjected to this stimulus and undisturbed controls.

#### Arousal Thresholds

Arousal thresholds were calculated using the Drosophila Arousal Tracking system (DART) as previously described^64,65^. Female flies were housed individually in 80mm glass tubes, and their activity was monitored using video tracking. 14 flies were used per genotype. Flies so housed were probed hourly for 24hr, with a train of vibrational stimuli of increasing strength from 0 to 1.2g. Each stimulus consisted of 5 pulses of 200ms, and was delivered in 0.24g increments 15 seconds apart. The arousal threshold for each fly was calculated as the weakest vibration intensity (g) required to elicit a response (walking at least half the length of the glass tube) in quiescent flies that had been inactive for least the preceding minute. The average arousal threshold across the day was then calculated for each strain.

#### Confinement

Individual female flies were collected pre-expansion and confined overnight in <7mm space as previously described. Following confinement, flies were placed in 65mm tubes for sleep recording.

#### Wing cut

Female flies were collected on the day they eclosed, and both wings were cut under C0_2_ anaesthesia after wings had expanded. Flies with cut wings, and their siblings with intact wings that had been subject to the same anaesthesia protocol, were then placed in 65mm glass tubes in DAM monitors. Sleep data is reported for second day post wing-cut i.e for 2 day old flies.

#### Wing glue

Wings of 3 day old female flies were glued with a small amount of UV activated glue (Bondic, ON) under CO_2_ anaesthesia and short (∼5s) UV light exposure. As a control a small amount glue was applied to the abdomen of siblings. Unglued flies were also subject to the same anaesthesia and UV curing light exposure. All flies were then housed in 65mm glass tubes and placed in DAM monitors.

### Immunohistochemistry

Whole flies were fixed in 4% paraformaldehyde (Electron Microscopy Sciences) in phosphate buffered saline (Sigma-Aldrich) + 0.3% Triton X-100 (PBST). Following fixation, the CNS was dissected in PBS, washed in PBST, and incubated in blocking solution (PBST + 5% normal goat serum) at 4° C overnight. The following day, brains and VNCs were incubated in primary antibodies (diluted in blocking solution) for 2 days at 4°C. Primary antibodies (and dilutions used) were: Chicken anti-GFP (1:1000, Abcam); Mouse monoclonal anti-GFP (1:100, Sigma, used in GRASP experiments referred to as anti-GRASP); Rabbit anti-Dsred (1:250, Takara Bio, used to label mcherry, tdtomato, RFP, etc.); rat anti HA (1:500, Sigma); mouse anti TH (1:500, Immunostar), mAb nc82 (1:400, DSHB); mouse anti V5 (1:400, Invitrogen). Following incubation in primary, brains were washed and incubated overnight in secondary antibody solution. Secondary antibodies used included: Goat anti-chicken Alexa Fluor 488 (1:400, Invitrogen), Goat anti-mouse Alexa Fluor 488 (1:400, Invitrogen), Goat anti-rabbit Alexa Fluor 568 (1:300, Invitrogen); Goat anti-rabbit Alexa Fluor 633 (1:300, Invitrogen); Goat anti-rat Alexa Fluor 568 (1:200, Invitrogen); Goat anti-mouse Alexa Fluor 568 (1:200, Invitrogen); Goat anti-mouse Alexa Fluor 633 (1:200, Invitrogen). Following incubation in secondary antibodies, brains were washed in PBST and mounted in Vectashield (Vector Labs). Images were obtained using an Olympus FV-1200 laser scanning confocal, using one of a UAPO 20X air, 40X water immersion, 63X water immersion, or with a Zeiss LSM 880 confocal microscope with a 40X oil objective. Confocal Z-stacks were acquired with sequential scanning frame by frame, to prevent bleedthrough across channels, and at a depth of 1.0µm or 0.5µm. Images were processed using Fiji or Imaris (Bitplane) software. Unless otherwise specified, 5-7 day old female flies were used for immunostaining experiments.

#### CaLexA measurements

CaLexA was expressed in R2 ellipsoid body neurons with R30G03 GAL4 or in VNC projection neurons with 31C06 GAL4. 3 day old CaLexA expressing female flies with cut wings and their siblings with intact wings were fixed at ZT0-1. CaLexA driven GFP signal in brains and VNCs was enhanced by staining with chicken anti-GFP. Images were acquired with a 40X water immersion objective at 1024 × 1024 pixels on an Olympus FV-1200 confocal microscope. Cut and intact groups were imaged using the same settings. Images were analysed using Fiji/ ImageJ. To measure fluorescent intensities, the sum of all pixels of a stack in a region of interest (ROI) was calculated. ROI intensities were corrected for background by measuring and subtracting background fluorescent intensity from a region adjacent to the ROI.

#### BRP measurements

STaR (synaptic tagging with recombination) was expressed with 31C06 Gal4 to label BRP puncta in VNC projection neurons. 3 day old 31c06 > STaR female flies with cut, curly and intact wings were fixed at ZT0-1. BRP puncta were visualised by immunostaining with an anti V5 antibody. Images were acquired with a 60X water immersion objective at 1024 × 1024 pixels on an Olympus FV-1200 confocal microscope. All groups were imaged with the same settings. 31C06 GAL4 labelled neurons with dendritic arborisations in the VNC, including in the wing neuromere, and axonal projections that exited the VNC forming two tracts along the sub-esophageal ganglion– a medial tract and a lateral tract. This nomenclature also reflected the position of these tracts in the VNC-brain connective. Both tracts terminated in the ventro-lateral protocerebrum (VLP) where they made extensive arborisations. Although the entire CNS was dissected and immunostained, we focused on the region of the brain containing the projections of the 31C06 wing-VLP projection neurons from the ventral SEG to the VLP. Anatomical experiments suggested that the 31C06 projection neurons are largely axonal in this region. Images were processed with Imaris software (Bitplane), which allows for visualisation and quantification of data in 3 dimensions. Images were analysed and quantified while being blinded to condition. Confocal z-stacks were carefully segmented by manual annotation of a ∼150µm z stack into 4 regions – region1 corresponding to the medial tract from the VNC to the VLP, region 2 the lateral tract, region3 the lateral (and anterior) VLP arborisations, and region 4 the dorsal (and posterior) arborisations of the two tracts in the VLP. Background intensity was automatically calculated for each region. Thresholding criteria for identifying BRP puncta were automatically generated, and occasionally manually adjusted. The number of BRP puncta and average staining intensity for each punctum was automatically generated.

#### Statistics

Statistical analyses were carried out in Systat software. Statistical comparisons were done with a Student’s t-test for comparisons between two groups, or ANOVA followed by Tukey’s post-hoc comparisons for tests involving multiple comparisons. Unless otherwise specified, the most commonly used statistical analysis was a one-way ANOVA for genotype / condition.

## Data Availability

All data is available in the main text or the supplementary materials; raw data are available from the corresponding author upon reasonable request.

## Acknowledgments

We thank L. Salkoff for critical input; T.Y.Lin and C.H. Lee for sharing reagents; H. Dierick, S. Dissel, M. Thimgan, and B. White for helpful discussions and comments on the manuscript; D. Oakley and M. Shih for technical input on image acquisition and analysis; L. Cao, Z. Koch, and D. Chan, for technical assistance. This work was supported by NIH grants 5R01NS051305-14 and 5R01NS076980-08 to PJS. The confocal facility is supported by NIH shared instrument grant S1OD21629-01A1.

## Author contributions

KM, BZ, and PJS conceived the project, designed and performed experiments, interpreted the data and wrote the manuscript.

---

**Extended Data Fig. 1:**
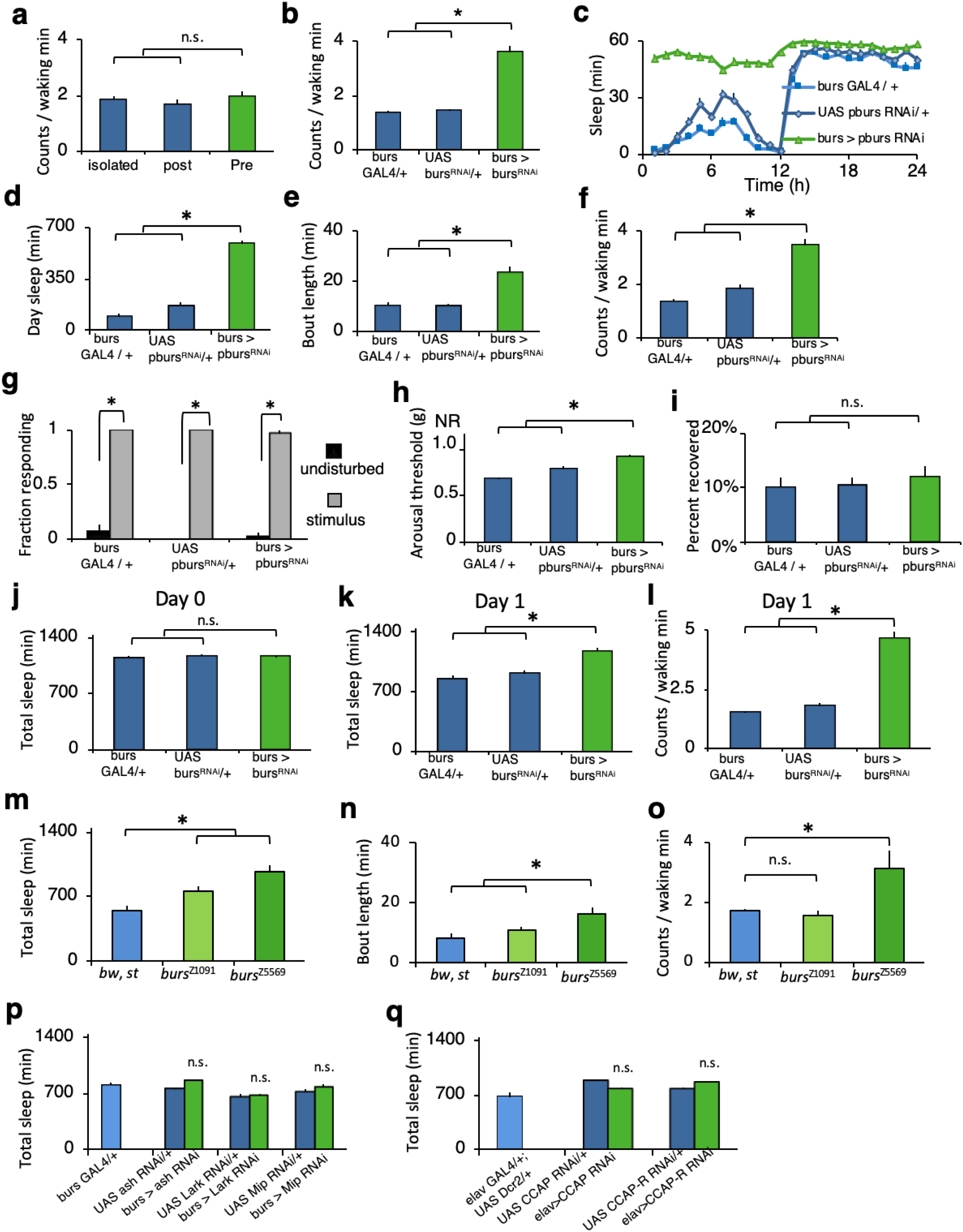
Extended characterisation of *burs* loss of function. **a**, Waking activity was not altered in flies confined pre-expansion (pre) compared to controls (n.s. p >0.05). The intensity of waking activity was elevated in 4-5 day old *burs GAL4* / + > *UAS burs RNAi* / + (**b**, p <0.001) flies compared to controls. **c**, *bursGAL4/+>UAS-pburs*^*RNAi*^*/+* slept more than parental controls (n=20-32 flies / genotype, Repeated measures ANOVA for Time X condition, p< 0.001). **d, e**, *bursGAL4/+>UAS-pburs*^*RNAi*^*/+* displayed increased daytime sleep and sleep bout duration compared to controls (*p< .01 Tukey correction). **f**, The intensity of waking activity was elevated in *bursGAL4/+>UAS-pburs*^*RNAi*^*/+* flies relative to controls. **g**, Sleep was rapidly reversible in response to a mechanical stimulus for all genotypes (n=25-30 flies / condition, *p< .01 Tukey correction). **h**, Sleep in *bursGAL4/+>UAS-pburs*^*RNAi*^*/+* flies was associated with increased arousal thresholds (n=14 flies per condition; *p< .01 Tukey correction). **i**, All genotypes displayed similar sleep rebound following 12 h of sleep deprivation (n=21-32 flies / condition). **j**, In contrast to confinement, *burs GAL4* / + > *UAS burs RNAi* / + flies did not alter sleep on the day flies eclosed relative to parental controls (day0, n.s. p>0.44 Tukey correction). **k**, Sleep was significantly elevated however, in *burs GAL4* / + > *UAS burs RNAi* / + flies on the day after eclosion (Day1, n=21 flies per genotype, * p< 0.001 Tukey correction). **l**, The intensity of waking activity was elevated in these flies on day 1 (* p< 0.001). **m**, Loss of function *burs* mutants exhibited increased sleep (* p < 0.05, n=16 flies per condition), increased sleep consolidation during the day (**n**,* p <0.05), and did not decrease waking activity (**o**, n.s. p>0.05, * p<0.05). **p**, Sleep of *burs GAL4* / + > *UAS ash RNAi* / +, *burs GAL4* / + > *UAS lark RNAi* / +, and *burs GAL4* / + > *UAS mip RNAi* / + flies was not altered compared to parental controls (n=16 flies / genotype, n.s. p> 0.44 Tukey correction).**q**, *elav GAL4* / + > *UAS CCAP RNAi* / + and *elav GAL4* / + > *UAS CCAPR RNAi* / + flies did not alter sleep relative to controls (n=16 flies / genotype, n.s. p>0.99 Tukey correction).

**Extended Data Fig. 2:**
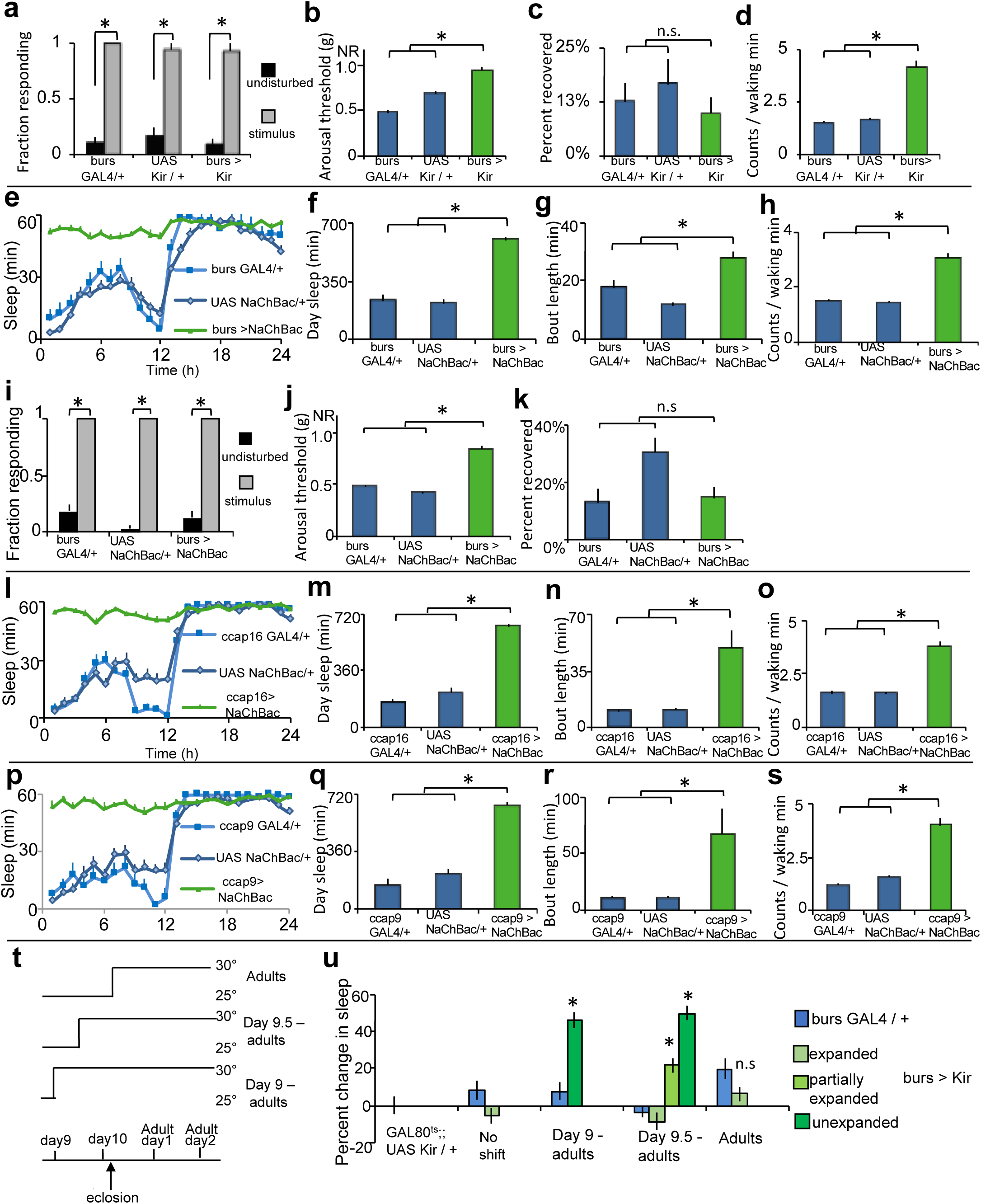
Manipulating the activity of *burs* neruons increases sleep. **a**, Sleep was rapidly reversible in response to a mechanical stimulus for all genotypes (n=26-30 flies / condition, *p< .01 Tukey correction). **b**, Sleep in *bursGAL4/+>UAS-Kir2.1/+* flies was associated with increased arousal thresholds (n=14 flies per condition; *p< .01 Tukey correction), **c**, All genotypes displayed similar sleep rebound following 12 h of sleep deprivation (n=20-30 flies / genotype). **d**, Waking activity was not decreased in *bursGAL4*>*Kir* flies relative to controls (* p<0.001). **e**, *burs GAL4* / + > *UAS NaChBac* / + flies increased sleep (n=24-30 flies / genotype, repeated measures ANOVA for time X genotype p< 0.001) particularly in the day (**f**, * p< 0.001), and increased daytime sleep consolidation (**g**, * p<0.01) relative to controls. **h**, Waking activity was elevated in *burs GAL4* / + > *UAS NaChBac* / + flies relative to controls (*, p <0.001). **i**, All genotypes were similarly awakened by a mechanical stimulus (n=30 flies / condition, * p< 0.001 Tukey correction). **j**, *burs GAL4* / + > *UAS NaChBac* / + flies exhibited higher arousal thresholds during the day (n=14 flies per condition, * p<0.001), and a similar level of rebound sleep following sleep deprivation to parental controls (**k**, n=27-31flies / genotype, p>0.08). **l** *CCAP-16 GAL4* / *+> UAS NaChBac / +* flies increased sleep relative to parental controls. (n=16 flies / genotype, repeated measures ANOVA for time X genotype,, * p < 0.001). **m**-**o**, *CCAP-16 GAL4* / *+> UAS NaChBac / +* flies displayed increased daytime sleep (* p< 0.001 Tukey correction), sleep bout length (* p < 0.01), and waking activity throughout the day (*, p< 0.001) relative to controls. **p**, *CCAP-9 GAL4* / *+> UAS NaChBac / +* flies increased sleep relative to parental controls. (n=16 flies / genotype, repeated measures ANOVA for time X genotype, * p < 0.001). **q**-**s**, *CCAP-9 GAL4* / *+> UAS NaChBac / +* flies displayed increased daytime sleep (* p< 0.001 Tukey correction), sleep bout length (* p < 0.05), and waking activity throughout the day (*, p< 0.001) relative to controls. **t**, *burs* neurons were inhibited by expressing the inward rectifying K+ channel Kir2.1 with *burs* GAL4. A temperature sensitive GAL8O, *GAL80*^*t*s^ was used to temporally restrict the activity of *burs GAL4*, and thus the inhibition of *burs* neurons to defined time windows (*burs GAL4* / + > *GAL80*^*ts*^ / + ; *UAS Kir* / +). Flies were reared at 25°C; in these conditions they eclose 10days after egg-laying (day10). Days 9-10 represent days after egg laying, Adult Day1&2 represent days after eclosion. Sleep data is shown for the second day post eclosion. **u**, Raising *burs GAL4* / + > *GAL80*^*ts*^ / + ; *UAS Kir* / + flies at the permissive temperature throughout development (No shift, n=31 flies / genotype) or shifting flies to the restrictive temperature post wing expansion (adults, n=27-30 flies/ genotype) had no effect on wing expansion or sleep. In contrast, shifting *burs GAL4* / + > *GAL80*^*ts*^ / + ; *UAS Kir* / +flies to the restrictive temperature one day prior to eclosion (Day 9-adults, n=28-30 flies / genotype, * p<0.001 Tukey correction) or ∼12 hours prior to eclosion (Day9.5-adults) blocked wing expansion, with the flies that emerged with wing defects (∼80% of progeny, n=40) exhibiting increased sleep relative to controls (n=32flies / genotype, * p <0.001). All manipulations blocked wing expansion.

**Extended Data Fig. 3:**
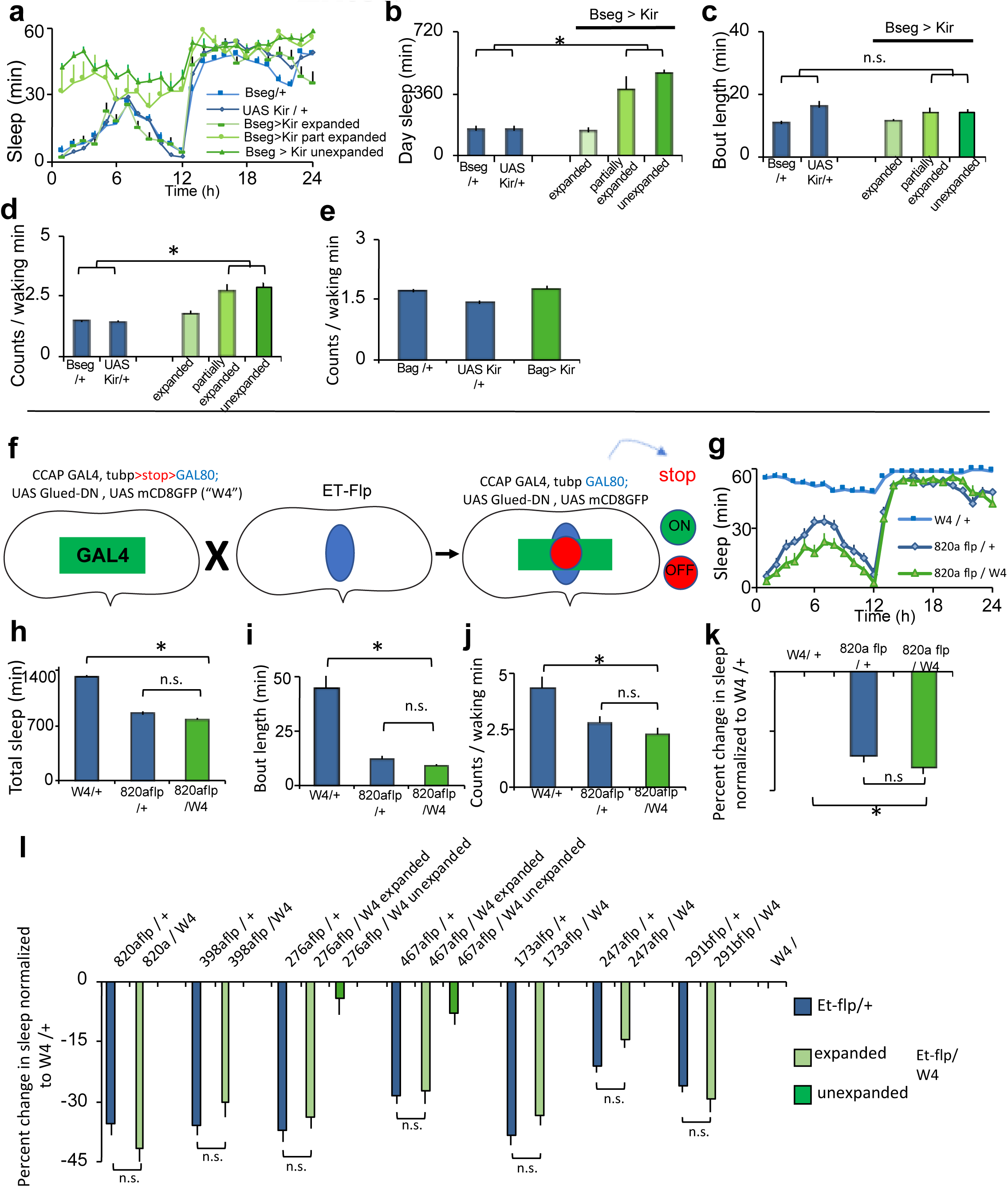
Intersectional strategy to define a minimal subset of burs/CCAP expressing neurons. **a** Inhibition of the Bseg partially blocked wing expansion with ∼60% of Bseg>Kir progeny displaying wing defects, and increased sleep (repeated measures ANOVA for time X genotype p< 0.001). Bseg > Kir flies with wing expansion defects increased sleep during the day (**b**, * p< 0.001) without changing sleep consolidation (**c**) and were more active while awake (**d**, * p <0.001). **e**, Waking activity was unchanged in Bag>Kir flies (n.s. p> 0.64). **f**, Summary of the FINGR method, CCAP GAL4 neurons are disrupted by expressing a Glued-DN transgene with CCAP GAL4 to disrupt the cytoskeleton and thereby neural function. A FRT flanked stop cassette blocks the expression of GAL80, permitting GAL4 activity (“W4 flies”). This manipulation blocks wing expansion and generates long-sleeping flies. The W4 flies were crossed to a number of enhancer-trap flipase lines (ET-flp). GAL80 is “flipped in” and GAL4 consequently repressed, in cells defined by the intersection of the expression pattern of the CCAP enhancer, and the enhancer driving the flipase. **g**, Representative sleep profile of an Et-flp line (820a flp) that suppressed the wing expansion defects seen in W4 flies (n=22-30 flies/ genotype). **h**, Sleep of *820a flp* / + > *W4* / + flies is indistinguishable from the *820a flp / +* parental control line (*, p< 0.001, n.s. p> 0.77). Similar changes were seen in day bouts (**i**, *, p< 0.001, n.s. p> 0.99) and waking activity (**j**, *, p< 0.001, n.s. p> 0.40). **k**, Sleep data in (**h**) expressed as a percentage of change in sleep amount normalised to the W4 / + parental control line. W4 flies are long-sleeping flies, so a ‘normal’ fly would be expected to exhibit a dramatic decrease in sleep relative to W4 flies. (**l**) Seven different ET-flp lines that were previously shown to suppress wing expansion defects when crossed to W4 flies, were obtained and crossed to W4 flies. Sleep is expressed as the percent change in total sleep relative to W4 flies. All of the progeny with expanded (normal) wings reduced sleep compared to the W4/+ parental control (p< 0.001), to levels statistically indistinguishable from their parental Et-flp / + controls (n.s. p> 0.20). In contrast, progeny with unexpanded wings had levels of sleep significantly higher than their Et-flp / + controls (p< 0.001), but indistinguishable from the W4 / + control (p>0.10). These Et-flp lines overlapped with CCAP GAL4 in the subesophageal ganglion. n=12-30 flies per condition.

**Extended Data Fig. 4:**
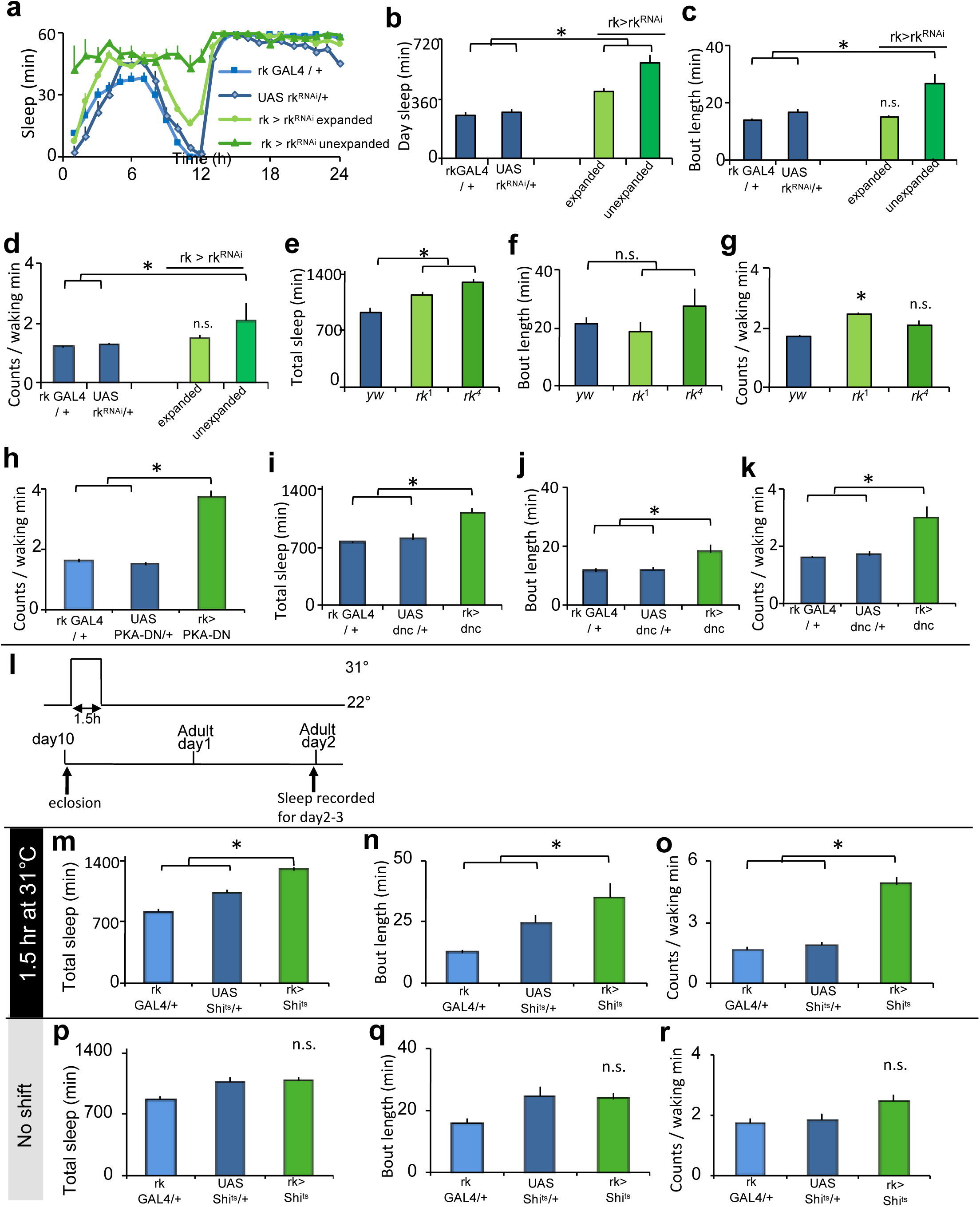
Disrupting *rickets* signalling blocks wing expansion and increases sleep. **a**, Knocking down *rk* with RNAi (*rk GAL4* / + > *UAS rk RNAi* / +) resulted in a partially penetrant effect on wing expansion (with ∼40% of progeny displaying wing defects). Two way repeated measures ANOVA detected a significant effect of genotype on time (n=16 flies / condition, p<0.001). **b**, Daytime sleep was elevated in *rk GAL4* / + > *UAS rk RNAi* / + flies with expanded wings and *rk GAL4* / + > *UAS rk RNAi* / + flies with unexpanded wings relative to controls (*, p< 0.05, Tukey correction). **c, d**, *rk GAL4* / + > *UAS rk RNAi* / + flies with unexpanded wings showed increased sleep consolidation during the day (* p<0.01, n.s. p> 0.8) and increased waking activity (* p < 0.01, n.s. p> 0.17) compared to controls. **e**, Sleep was elevated in two different loss of function *rickets* (*rk*) mutants – *rk*^1^ & *rk*^4^ compared to controls (n=12-14 flies / genotype *, p< 0.01 Tukey correction). **f**, Daytime sleep consolidation was not altered in either mutant relative to controls (n.s. p>0.28). **g**, Waking activity was not decreased in either mutant relative to controls (* p <0.05, n.s. p > 0.17). **h**, The intensity of waking activity was elevated in rk > PKA-DN flies compared to controls (*, p <0.001). **i**, Sleep was increased in *rk GAL4* / + > *UAS dnc* / + flies (n=8) compared to controls (n=12-16 flies / genotype *, p <0.01 Tukey correction). **j, k**, *rk GAL4* / + > *UAS dnc* / + flies displayed sleep consolidation during the day (* p <0.05) and increased waking activity (* p < 0.001) compared to controls. **l**, A temperature sensitive dynamin (UAS Shi^ts^) was used to inducibly block chemical transmission from *rk*GAL4 expressing neurons. *rk GAL4* / + > *UAS Shi*^*ts*^ / + flies and parental controls were shifted to the restrictive temperature of 31° for 1.5 hrs just after eclosion. This manipulation was sufficient to block wing expansion. Flies eclose ∼10 days after egg-laying in these conditions (day10). Sleep data is shown for the second day after eclosion (Adult day2-3) for flies subjected to the temperature shift (**m**-**o**, n=12-16 flies / genotype) and sibling flies maintained at the permissive temperature (**p**-**r**, n=14-16 flies / genotype). Maintaining *rk GAL4* / + > *UAS Shi*^*ts*^ / + flies at the permissive temperature did not affect wing expansion.(**m**-**o**) *rk GAL4* / + > *UAS Shi*^*ts*^ / + flies subjected to the temperature shift increased sleep (* p< 0.001 Tukey correction), sleep consolidation during the day (* p<0.01), and intensity of waking activity * p<0.001)) compared to parental controls. (**p**-**r**) In contrast, sleep architecture of *rk GAL4* / + > *UAS Shi*^*ts*^ / + flies maintained at the permissive temperature was not altered relative to controls.

**Extended Data Fig. 5:**
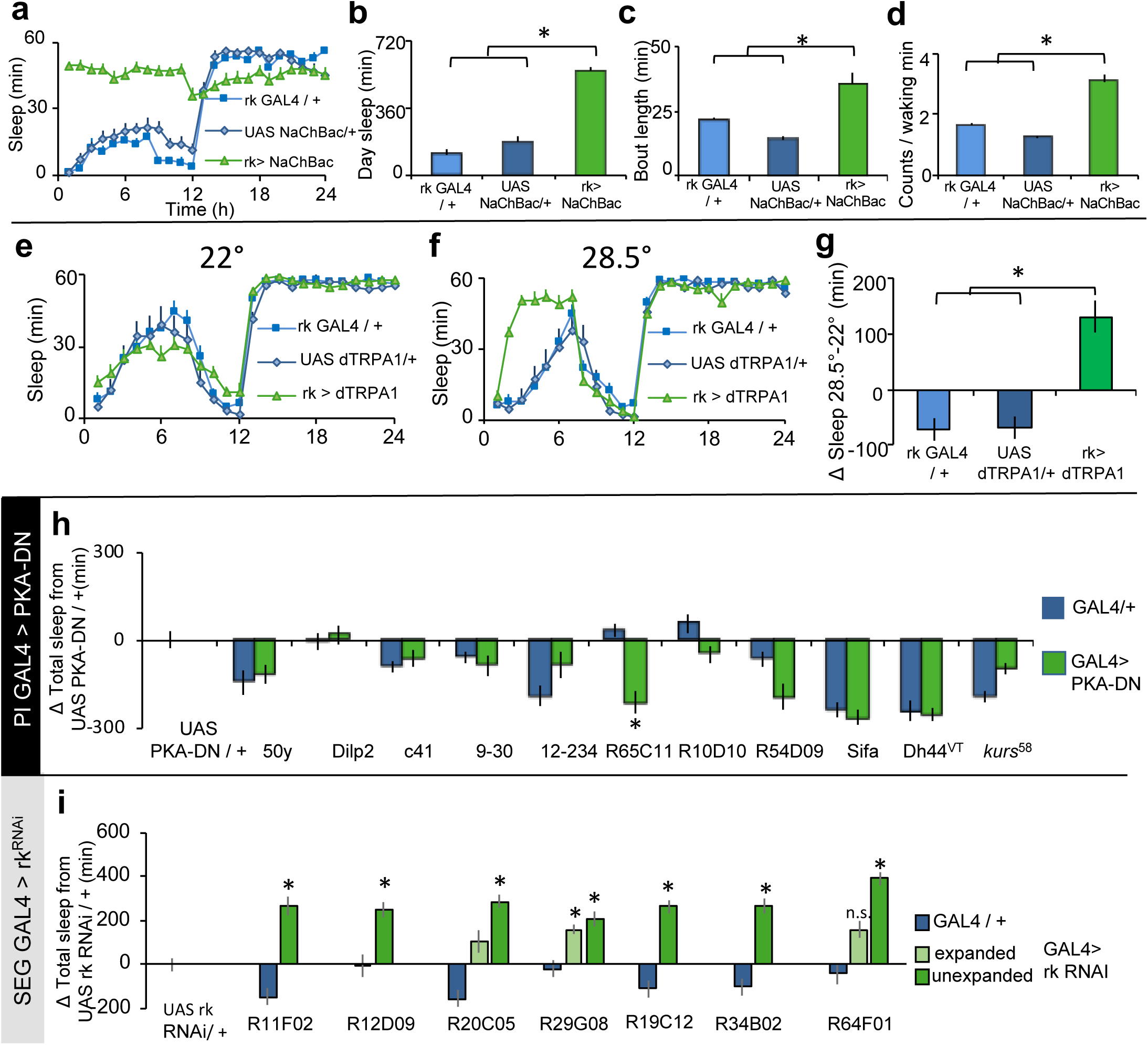
Manipulating *rk* neurons increases sleep. **a**-**d**, Chronic activation of *rk GAL4* expressing neurons with NaChBac blocks wing expansion and increases sleep **e**-**g**, Acute activation *rk GAL4* expressing neurons with dTRPA1 in 4-5day old adult flies increases sleep. **a**, *rk GAL4* / + > *UAS NaChBac* / + flies increased sleep relative to controls (n=18-30 flies / genotype, Two-way repeated measures ANOVA for time X genotype (p<0.001). **b**-**d**, Daytime sleep amount (* p< 0.001), sleep consolidation (* p< 0.001), and waking activity (* p< 0.001) were all elevated in *rk GAL4* / + > *UAS NaChBac* / + flies compared to both controls. **e**, Sleep plot of *rk GAL4* / +> *UAS dTRPA1* / + flies and parental controls at the permissive temperature (23°) on the baseline day (n-12-16 flies / genotype) **f**, Sleep plot of *rk GAL4* / +> *UAS dTRPA1* / + flies and parental controls for the day of the temperature shift. *rk* neurons were acutely activated on the day following the baseline day, by shifting flies to the restrictive temperature (28.5°) for 6 hrs (orange box). **g**, *rk GAL4* / +> *UAS dTRPA1* / + flies acutely increased sleep for the 6 hours of the temperature shift compared to parental controls (*, p < 0.001 Tukey correction). Sleep is quantified as 6 hr totals of sleep at restrictive temperature – 6 hr totals at permissive temperature for each genotype. **h**, We noticed that the *rk GAL4* transgene drove expression in neurons of the pars intercerebralis (PI), a known sleep regulatory centre. To investigate the potential role of the PI in mediating the effects of *rk* signalling on wing expansion and sleep we expressed a dominant negative PKA in subsets of PI neurons with different drivers. One of the lines decreased (R65C11) decreased sleep relative to controls (*, p<0.005 Tukey correction). None of the other lines significantly altered sleep. Importantly, in contrast to inhibiting PKA with *rk GAL4* or *R64F01 GAL4*, none of the lines increased sleep or blocked wing expansion (n=12-16 flies / genotype). **i**, Since PKA is known to be downstream of many G protein coupled receptors (GPCRs), we did a secondary screen (with UAS *rk* RNAi) of the GAL4 lines that were hits from the PKA-DN screen. Sleep is shown as change in sleep normalised to the *UAS rk RNAi* / + parental control line. Driving rk RNAi with seven different GAL4 lines disrupted wing expansion and increased sleep (*, P<0.01 Tukey correction). These lines do not appear to overlap in their expression patterns, but instead express in different subsets of neurons in the subesophageal ganglion n=16 flies per genotype. PKA – protein kinase A.

**Extended Data Fig. 6:**
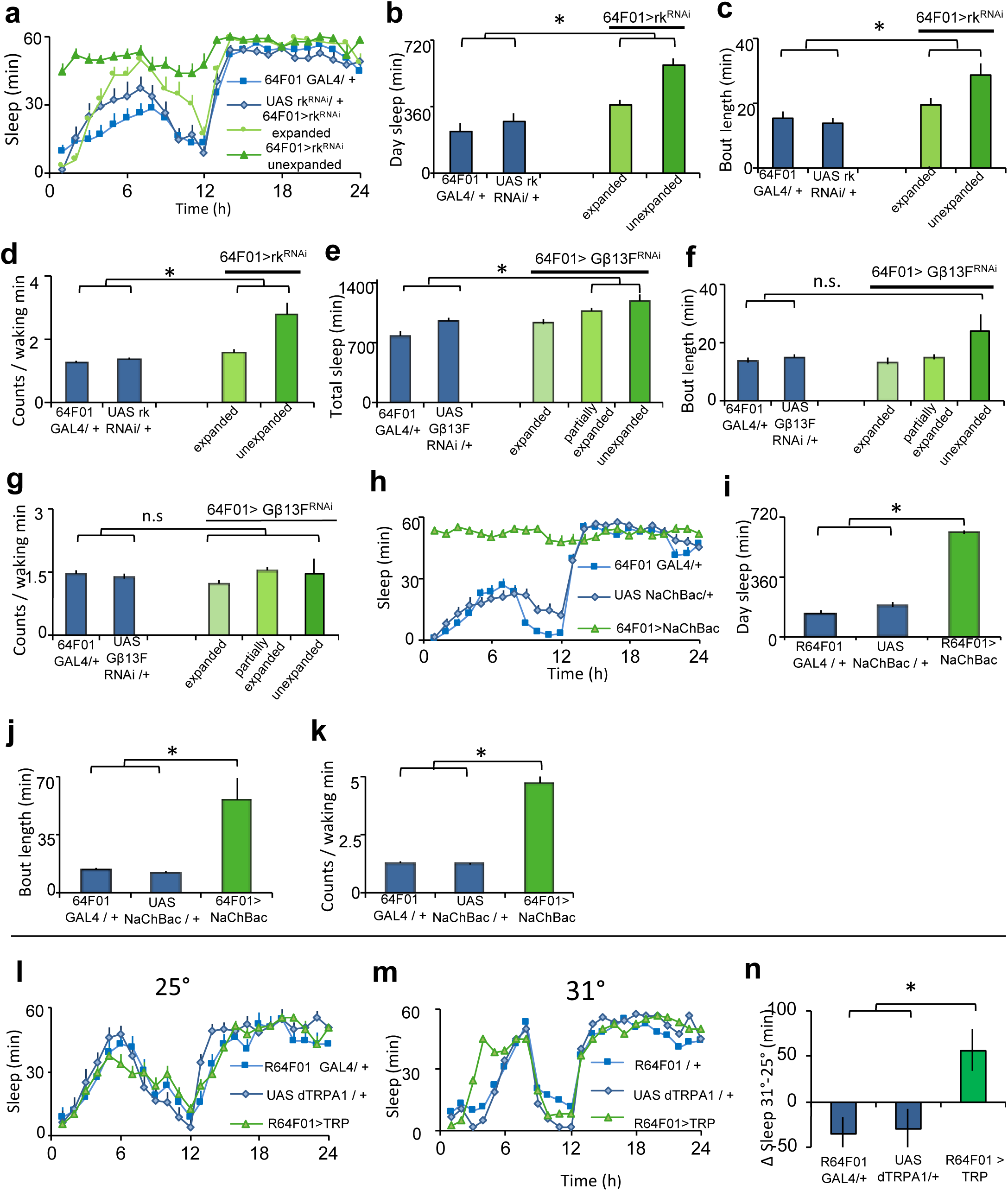
Manipulating components of *rk* signalling in, and activating R64F01 GAL4 neurons increases sleep. **a**-**d**, Driving *rk* RNAi with R64F01 GAL4 had a partially penetrant effect on wing expansion with ∼48% of *R64F01 GAL4 / +* > *UAS rk RNAi* / + flies exhibiting wing expansion defects. **a**, Sleep plot of *R64F01 GAL4 / +* > *UAS rk RNAi* / + flies and parental controls. Two-way repeated measures ANOVA detected a significant genotype X time interaction (p<0.001). **b**-**d**, *R64F01 GAL4 / +* > *UAS rk RNAi* / + flies with unexpanded wings (n=8) increased daytime sleep (* p <0.01 n.s. p > 0.10), sleep consolidation (* p< 0.01), and waking activity (* p<0.001) compared to controls (n=13-16 flies per genotype). **e-g**, The *rk* receptor is known to signal through the Gβ subunit Gβ13F. Knocking down Gβ13F with RNAi in *R64F01 GAL4* neurons (*R64F01 GAL4* / + > *UAS Gβ13F RNAi /* +) resulted in a partially penetrant effect on wing expansion with ∼68% of progeny displaying wing defects (n=53).**e**, *R64F01 GAL4* / + > *UAS Gβ13F RNAi /* + flies with wing expansion defects increased sleep compared to parental controls (n=32 flies / genotype, * p<0.001 Tukey correction). **f**-**g**, Sleep consolidation and waking activity were not altered in *R64F01 GAL4* / + > *UAS Gβ13F RNAi /* + flies relative to controls. (**h**-**k**) Chronic activation of *R64F01 GAL4* expressing neurons with NaChBac blocks wing expansion and increases sleep (**l**-**n**) Acute activation of *R64F01 GAL4* expressing neurons with dTRPA1 in 4-5day old adult flies increases sleep. **h**, *R64F01 GAL4* / + > *UAS NaChBac* / + flies increased sleep relative to controls (n=20-30 flies / genotype, two-way repeated measures ANOVA for time X genotype p<0.001). **i**-**k**, Daytime sleep amount (* p< 0.001 Tukey correction), sleep consolidation (* p< 0.001), and waking activity (* p< 0.001) were all elevated in *R64F01 GAL4* / + > *UAS NaChBac* / + flies compared to both controls. **l**, Sleep plot of *rk GAL4* / +> *UAS dTRPA1* / + flies and parental controls at the permissive temperature (23°) on the baseline day (n-12-16 flies / genotype). **m**, Sleep plot of *R64F01 GAL4* / +> *UAS dTRPA1* / + flies and parental controls for the day of the temperature shift (n=14-16 flies / genotype). *R64F01 GAL4* neurons were acutely activated on the day following the baseline day, by shifting flies to the restrictive temperature (31°) for 6 hours (orange box). **n**, *R64F01 GAL4* / +> *UAS dTRPA1* / + flies acutely increased sleep for the 6 hours of the temperature shift compared to parental controls (*, p < 0.05 Tukey correction). Sleep is quantified as 6 hr totals of sleep at restrictive temperature – 6 hr totals at permissive temperature for each genotype.

**Extended Data Fig. 7:**
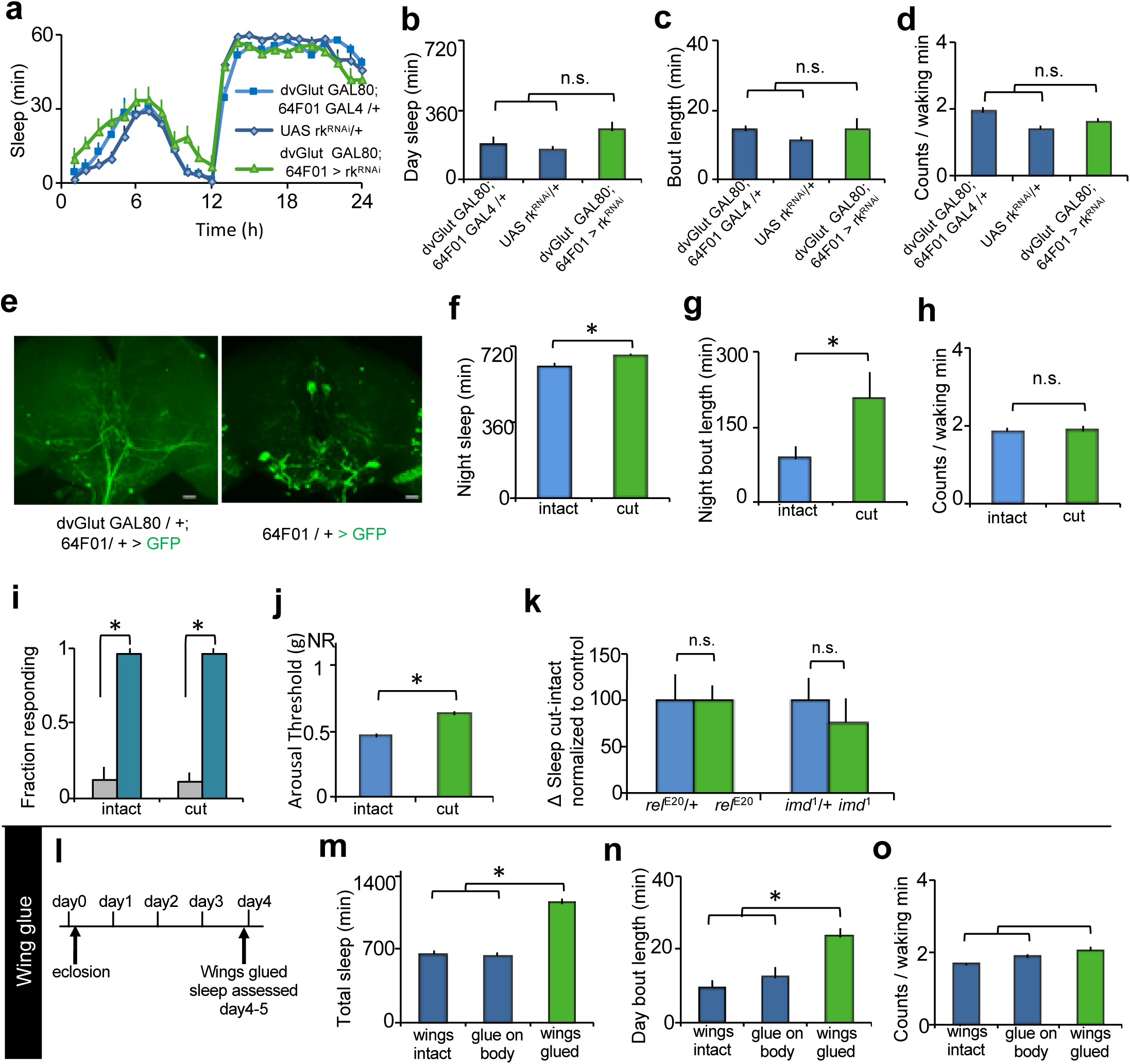
*rk* is required in putative glutamatergic neurons, and detailed characterization of wing-disruption induced sleep. **a**-**d**, *rk* is likely required in glutamatergic R64F01 neurons. *dvGlut GAL80* was used to restrict *R64F01 GAL4* mediated expression to non-glutamatergic neurons. **a**, Sleep plot of *dvGlut GAL80; R64F01 GAL4 / +* > *UAS rk RNAi* / + flies and parental controls (n=15-16 flies/ genotype). This manipulation did not affect wing expansion. **b-d**, *dvGlut GAL80; R64F01 GAL4 / +* > *UAS rk RNAi* / + flies did not change sleep amount (n.s. p >0.3), consolidation (n.s. p >0.88), or waking activity (n.s. p >0.7). **e**, Comparison of GFP expression driven by *dvGlut GAL80*; *64F01 GAL4* with that of sibling controls lacking the *dvGlut GAL80* transgene reveals that *dvGlut GAL80* suppresses expression in most *R64F01* GAL4 neurons, including in the prominent subesophageal ganglion neurons labelled by this line (Green, GFP). Maximal intensity confocal projections, scale bar 20μm. **f**-**h**, Wing cut increased sleep amount (* p <0.05 Student’s t-test) and consolidation (* p < 0.05 Student’s t-test) at night, without changing waking activity (n.s. p >0.88 Student’s t-test) compared to intact wing controls. **i**, Both flies with cut wings and their sibling controls with cut wings were equally aroused by a mechanical stimulus delivered at ZT15, on the second day following wing cut (*, p<0.001 Tukey correction) **j**, Flies with cut wings displayed a greater arousal threshold during the day compared to their siblings with intact wings (* p < 0.001 Student’s t-test). **k**, Mutations in immune system genes did not impair the ability of wing cut to induce sleep. Wing-cut dependent sleep is quantified for flies homozygous mutant for immune deficiency (*imd*^1^) gene and the NFκB *relish* (*rel*^e20^) as the change in sleep (cut-intact) normalized to the appropriate heterozygous allele (n.s. p >0.1 Student’s t-test). **l**, Protocol for wing glue. Wings of 3 day old adult flies were glued, and sleep assessed on the day following wing glue. **m**-**o**, Flies with wings glued slept more (n=19-36 flies/ condition, * p <0.001 Tukey correction), with more consolidated bouts during the day (* p <0.001) without changing activity while awake (n.s. p >0.06) compared to controls with intact wings and with an equivalent amount of glue applied to the abdomen (glue on body).

**Extended Data Fig. 8:**
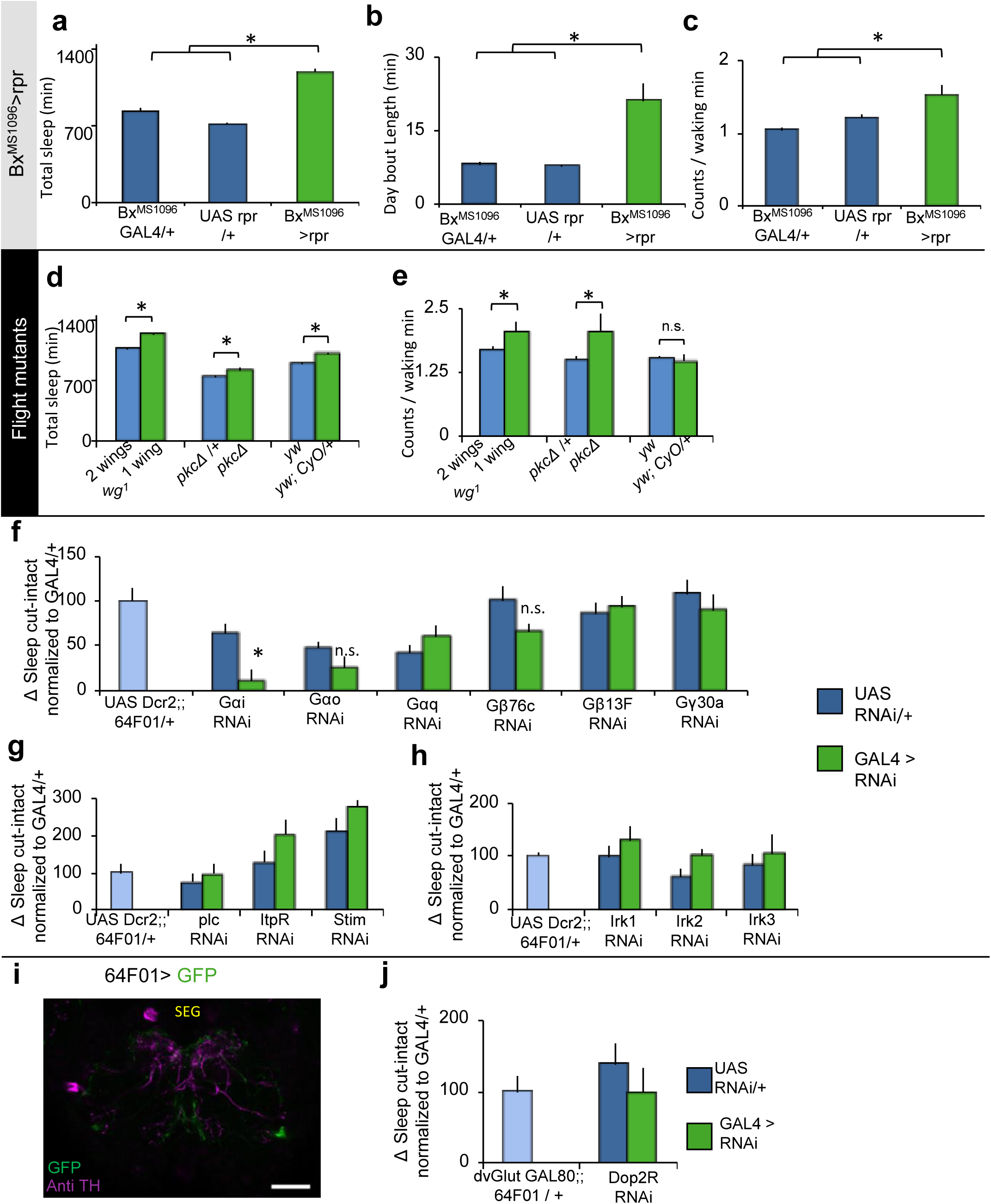
Additional characterisation of flight mutants and neuromodulation in wing-cut induced sleep. **a**-**c**, The cell-death activator reaper (rpr) was expressed in the wing disc with Bx^MS1096^ GAL4 driver. This manipulation generated flies with unexpanded wings. Importantly since the Bx^MS1096^ GAL4 driver expresses in the wing epithelial primordium, not in central brain wing expansion circuits, this manipulation is a way to impair wing expansion independent of *bursicon / rickets* signalling in the brain. *Bx*^*MS1096*^ *GAL4 / +* > *UAS rpr* / + flies increased sleep (**a**, n=14-28 flies / genotype * p <0.001), sleep consolidation (**b**, * p <0.001) and were more active while awake compared to parental controls (**c**, * p < 0.01). **d**, Sleep of three different mutations that are known to impair flight was evaluated compared to appropriate controls. The canonical *wingless* mutation (*wg*^1^) mutation has a partially penetrant effect on wing development with some homozygous mutant *wg*^1^ flies emerging with two wings and some with one wing. *wg*^1^ mutant flies with one wing slept more than their siblings with two wings (n=12-20 flies / genotype * p <0.001 Student’s t test). Similarly, flies carrying the dominant marker CyO slept more than their wild-type siblings (n=19-26 flies/genotype, *, p<0.05 Student’s t test). Flies homozygous mutant for protein kinase c Δ (pkcΔ) also slept more than their heterozygous controls (n=7-23 flies / genotype, * p<0.001 Student’s t-test). **e**, The intensity of waking activity was not decreased in any of the mutants relative to controls (*, p < 0.05, n.s. p>0.05). **f**, Different G protein subunits were knocked down in R64F01 neurons with RNAi. The extent of wing-cut induced sleep was significantly reduced in *UAS Dcr2;; R64F01 GAL4 /* + > *UAS Gαi RNAi* / + flies relative to controls (n=24-52 flies / condition, *, p<0.05, n.s. p > 0.10). **g**, Signalling through the Store Operated Calcium Entry (SOCE) in dispensable in R64F01 neurons for wing-cut mediated sleep increase. The extent of wing-cut induced sleep was not different in *UAS Dcr2;; R64F01 GAL4 /* + > *UAS plc RNAi* / +, *UAS Dcr2;; R64F01 GAL4 /* + > *UAS ItpR RNAi* / +, or *UAS Dcr2;; R64F01 GAL4 /* + > *UAS Stim RNAi* / + flies relative to controls (n=16-32 flies / condition, n.s. p >0.54). **h**, Signalling through inward rectifying potassium channels is dispensable in R64F01 neurons for wing cut mediate sleep. The extent of wing-cut induced sleep was not different in *UAS Dcr2;; R64F01 GAL4 /* + > *UAS Irk1 RNAi* / +, *UAS Dcr2;; R64F01 GAL4 /* + > *UAS Irk2 RNAi* / +, or *UAS Dcr2;; R64F01 GAL4 /* + > *UAS Irk3 RNAi* / + flies relative to controls (n=26-32 flies / condition, n.s. p >0.98). **i**, *R64F01 GAL4 / +* > *UAS mCD8GFP / +* neural processes were detected in close proximity to dopaminergic neural processes (labelled by anti-TH) in the SEG. **j**, The extent of wing-cut induced sleep was not different *in dvGlut GAL80; 64F01 GAL4* / + > *UAS Dop2R RNAi* / + flies relative to controls (n=21-30 flies / genotype n.s. p >0.15). **i**, Single confocal slice, green – GFP, magenta – anti TH (tyrosine hydroxylase). Scale bar 20μm, SEG-subesophageal ganglion.

**Extended Data Fig. 9:**
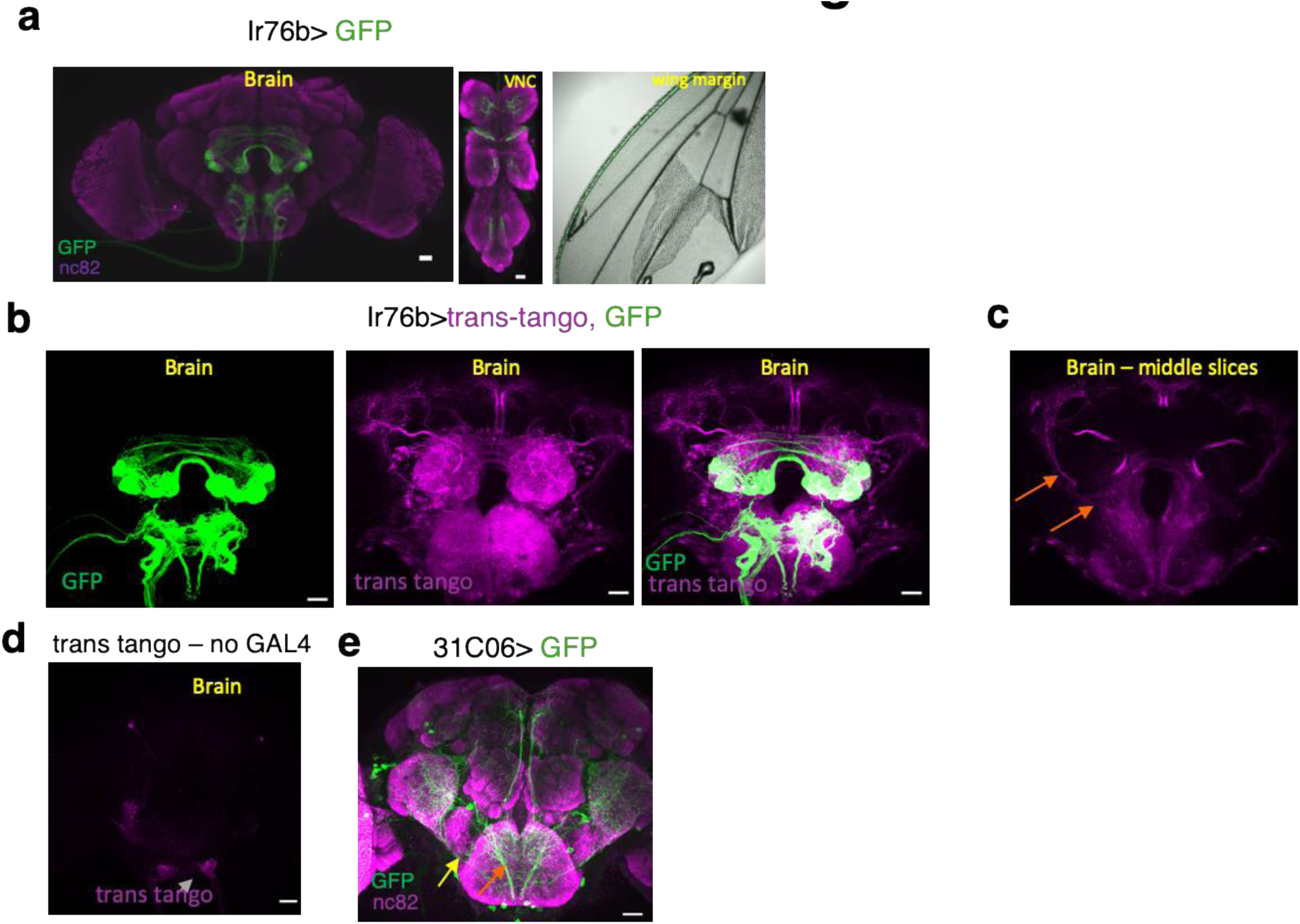
Additional anatomical characterisation related to Fig. 5. **a**, Like *Ir52a GAL4* (Fig 5b), GFP driven by *Ir76b GAL4* (green) also labelled sensory neurons in the wing (right), and sensory afferents from legs into the VNC (middle). In addition, *Ir76b GAL4* also drove expression in classes of olfactory sensory neurons and putative gustatory neurons in the labellum that project into the brain (left). CNS tissues were counterstained with nc82, a neuropil marker (magenta). **b**, Consistent with the expression of *Ir76b GAL4* in many different classes of sensory neurons (green, left), *Ir76b GAL4 > trans-tango* (magenta, middle) labelled a broad population of second-order neurons, with strong labelling in the SEG and antennal lobes, including olfactory projection neurons. **c**, A sub-stack of optical slices in (**b**), revealed I*r76b GAL4* > *trans-tango* expression in a projection neuron tract that resembled the medial tract seen in Fig. 5e (orange arrows). **d**, As a negative control, trans-tango components were crossed to a control yw strain, faint signal was detected in the ventral medial SEG (grey arrows). **e**, 3*1C06 GAL4* / + > *GFP* / + labelled projection neuronal tracts that connect the VNC to higher order brain centres along a medial (orange arrow) and a lateral tract (yellow arrow) with extensive arborization in the SEG and VLP Green – GFP, magenta – nc82. **b**-**e**, Images are maximal intensity z-projections of confocal stacks. Scale bar = 20μm. VLP – ventro-lateral protocerebrum, SEG – subesophageal ganglion.

**Extended Data Fig. 10:**
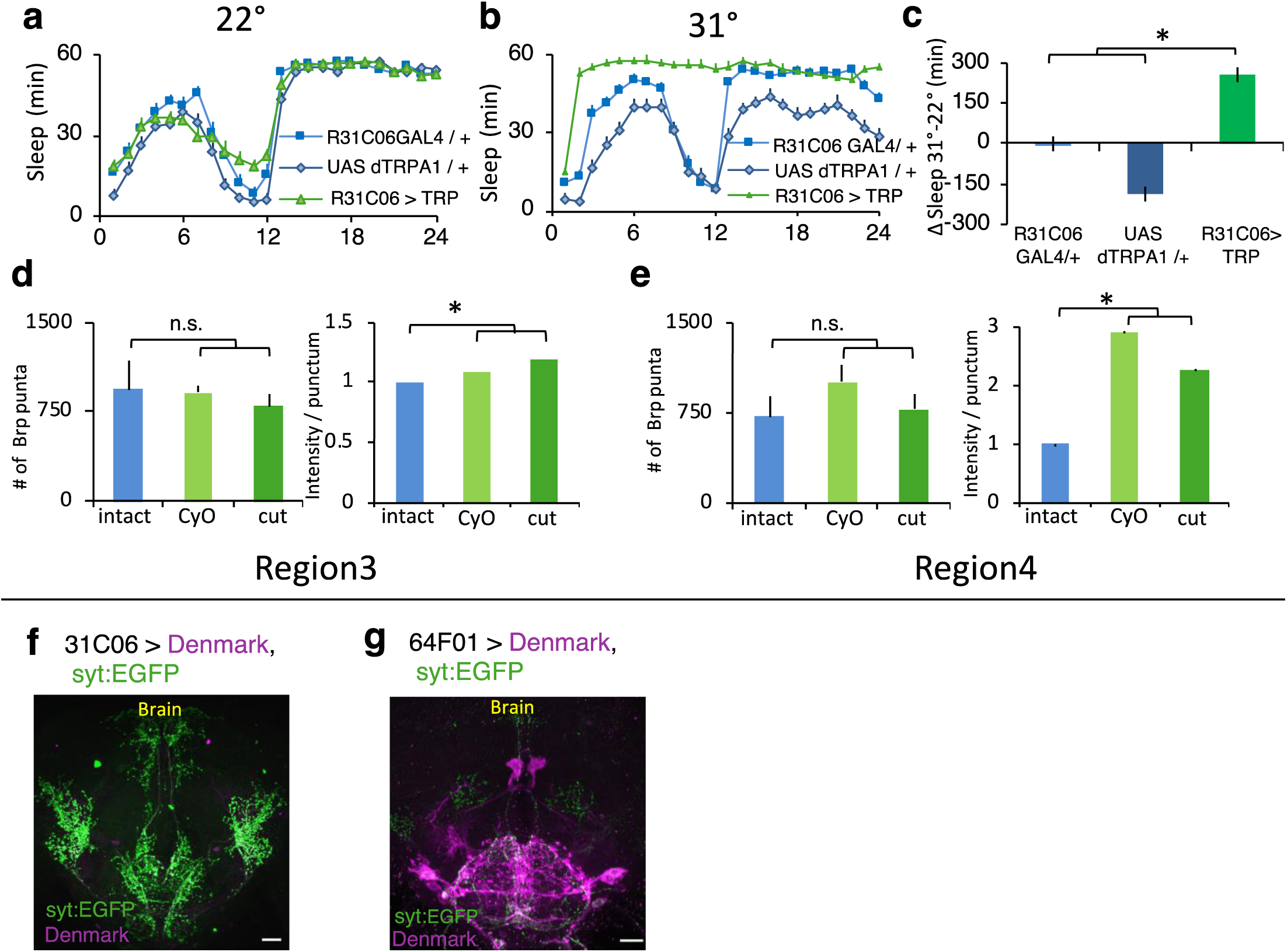
Activation of *R31C06 GAL4* neurons increases sleep, and additional anatomical characterization related to Fig. 6 **a**, Sleep plot of *R31C06 GAL4* / +> *UAS dTRPA1* / + flies and parental controls at the permissive temperature (22°) on the baseline day (n-20-30 flies / genotype) **b**, Sleep plot of *R31C06 GAL4* / +> *UAS dTRPA1* / + flies and parental controls for the day of the temperature shift. *R31C06 GAL4* neurons were acutely activated on the day following the baseline day, by shifting flies to the restrictive temperature (31°) for 24 hrs. **c**, *R31C06 GAL4* / +> *UAS dTRPA1* / + flies acutely increased sleep compared to parental controls (*, p < 0.001 Tukey correction). Sleep is quantified as total sleep at restrictive temperature – total sleep at permissive temperature for each genotype. **d**, In region 3, the intensity of BRP puncta was increased in cut *R31C06GAL4/+>STaR* flies and *R31C06GAL4/+>CyO/+;STaR* flies compared to intact controls. (*p <0.001, Tukey correction). The number of puncta was not altered, however. **e**, The intensity of BRP puncta was also increased in region 4 in cut *R31C06GAL4/+>STaR* flies and *R31C06GAL4/+>CyO/+;STaR* flies compared to intact controls without changing the number of BRP puncta (*p <0.001, Tukey correction). **f, g**, *31C06GAL4/+>UAS-Denmark, UAS syt:EGFP/+* and *R64F01GAL4/+>UAS-Denmark*,,*UAS syt:EGFP/+* staining patterns in the brain. *UAS-Denmark* (magenta) labels dendrites, syt:EGFP (green) labels presynaptic sites.

